# Coordinated dysregulation of modular gene activity in human neuropathologies

**DOI:** 10.64898/2026.08.25.747130

**Authors:** Gugene Kang, Michael C. Oldham

**Affiliations:** Department of Neurological Surgery, University of California, San Francisco, USA; Weill Institute for Neurosciences, University of California, San Francisco, USA

## Abstract

Understanding which genes are reproducibly dysregulated in which cell types is foundational knowledge for efforts to slow or reverse pathologies. For neuropathologies, such efforts rely primarily on differential expression analysis of single-nucleus RNA-seq (snRNA-seq) data. However, this strategy suffers from experimental and statistical challenges that limit marker gene reproducibility. We describe a novel strategy called Covariation Projection Analysis (**CoPA**) that combines the power of bulk sampling with the precision of single-cell methods. By ‘projecting’ bulk gene coexpression modules onto pseudobulked snRNA-seq cell types, CoPA reveals the cellular origins of highly reproducible genomic programs and their relative importance among cell types. By comparing CoPA projection patterns between normal and pathological human brain samples using **d**ifferential CoPA (**dCoPA**), we identify gene coexpression modules that are uniformly and reproducibly dysregulated in specific neocortical cell types in Alzheimer’s disease or schizophrenia. We share our findings through a novel web application called CoPA Cabana (https://oldhamlab.shinyapps.io/copacabana/).

## INTRODUCTION

Understanding how gene activity differs among neurobiological cell types in health and disease is a major goal of neuroscientific research. Today, most studies of normal and pathological human brain samples address this problem through differential expression (DE) analysis of cell types identified through single-nucleus RNA-seq (snRNA-seq). Although conceptually straightforward, snRNA-seq requires many experimental steps and analysis choices, each of which can introduce bias, inflate type I and II errors, and impair the reproducibility of marker genes. Well-known sources of experimental bias and error include loss of cytosolic mRNA, buffering of stochastic transcription by the nucleus^1,2^, variable capture efficiency of nuclei and transcripts^3–5^, multiplets^6–9^, contamination with ambient RNA^10–13^, and artifacts from cDNA amplification^14^. Importantly, these factors are much more likely to distort marker gene identification than cell type identification, since the former relies on a single noisy gene expression vector while the latter relies on thousands. Furthermore, investigator choices can dramatically alter the results of DE analysis. For example, the choice of DE algorithm – and even the software package version – can produce very different lists of marker genes from the same underlying data^15^. Combined with concerns about pseudoreplication bias^16–18^ and statistical power – particularly for low-expressed genes that dominate most snRNA-seq datasets^19,20^ – it is important to evaluate the reproducibility of marker genes identified by snRNA-seq and explore alternative strategies for studying pathological gene activity.

Genes do not function in isolation, but rather as members of coordinated genomic programs that support distinct cellular functions. We and others have shown that genome-wide coexpression analysis of bulk tissue samples is a powerful approach for revealing highly reproducible programs (or ‘modules’) of genomic activity^21–28^, since variation in the cellular composition of bulk tissue samples inevitably drives covariation of optimal markers for cell types and states at scale. Because one bulk sample may represent several million cells, large bulk datasets often represent *billions* of cells, providing enormous statistical power to survey the population structure of genomic activity while avoiding most sources of experimental bias and error that impact snRNA-seq studies. On the other hand, the cellular origins of most bulk coexpression modules are unknown. The complementary strengths and weaknesses of bulk and snRNA-seq data suggest their integration might clarify differences in genomic activity among neurobiological cell types in health and disease.

Here we introduce a novel approach called **Co**variation **P**rojection **A**nalysis (**CoPA**) that combines the power of bulk coexpression with the precision of single-cell analysis to reveal the cellular origins of highly reproducible modules of genomic activity. We focus on adult human neocortex and its 24 major cell types (or subclasses), which have been identified and validated as distinct transcriptional clusters in multiple snRNA-seq studies^29–32^. Using CoPA, we show that the overwhelming majority of gene coexpression modules in adult human neocortex are not cell-type-specific, but that subtle differences in the expression levels of coexpressed genes among cell types are nevertheless extremely reproducible. By comparing module projection patterns between snRNA-seq datasets derived from normal and pathological human brain samples using **d**ifferential CoPA (**dCoPA**), we identify gene coexpression modules that are uniformly and reproducibly dysregulated in specific neocortical cell types from patients with Alzheimer’s disease or schizophrenia. We share our results through a novel web application called CoPA Cabana (https://oldhamlab.shinyapps.io/copacabana/), which also enables CoPA for genome-wide coexpression networks in OMICON (https://theomicon.ucsf.edu), which are derived from standardized analysis of 116 datasets representing >17K normal and neoplastic human brain samples (see companion study by Eliscu et al.).

## RESULTS

### Human neocortical subclass markers vary across snRNA-seq datasets and DE methods

We first examined the reproducibility of human neocortical subclass markers identified by DE analysis of snRNA-seq datasets. We focused on two large studies^29,30^ that surveyed 253,177 nuclei from the dorsal frontal cortex (DFC) and 273,971 nuclei from the middle temporal gyrus (MTG) of 12 neurotypical adult donors using 10x Chromium v3 (Cv3; **Table S1**). Both studies annotated cell types using the Brain Initiative Cell Census Network (BICCN) reference taxonomy, which defines 24 subclasses and ∼130-150 ‘supertypes’^32^. Because subclasses are more clearly defined than supertypes and comprise the neocortex in more reproducible proportions (**Fig. S1**), we focused most analysis at this level. To avoid pseudoreplication bias and mitigate false discoveries^16–18^, we summed expression levels for each gene over all subclass nuclei from each donor prior to donor-level DE analysis with edgeR^33^ glmLRT or DESeq2^34^ (test = ‘LRT’), which were among the top performing methods in a benchmark study of DE analysis for single-cell datasets^18^. We compared each subclass to all other subclasses and defined marker genes as those that were significantly upregulated (FDR < .05) and unique (i.e., significant for only one subclass). We evaluated the reproducibility of marker genes identified by edgeR or DESeq2 in different datasets and brain regions (**Fig. 1, Fig. S2, Table S2-S5**). Over all subclasses and both DFC datasets, on average only 33.7% of markers were identified by both methods (**Fig. 1a,b**). In total, we identified only 110 genes as reproducible subclass markers in both datasets using both methods (**Fig. 1c**), and 79% of these (n=87) were non-neuronal (**Fig. 1d**). No neuronal subclass had more than five reproducible markers, and nine neuronal subclasses had none (**Fig. 1d**). To explore the basis for these findings, we formed subclass metacells from each dataset by averaging gene expression for all subclass nuclei, then clustered metacells by genome-wide dissimilarity (1-cor). This analysis revealed that most neuronal subclasses were extremely similar, with average genome-wide correlations of 0.94–0.95 among excitatory subclasses, 0.91–0.93 among inhibitory subclasses, and 0.9–0.91 among all neuronal subclasses; in contrast, non-neuronal subclasses were much more heterogeneous (**Fig. 1e,f**). Similar results were observed in MTG (**Fig. S2**). These findings underscore challenges with DE analysis of snRNA-seq data, consistent with previous reports^35–37^.

**Fig. 1.**
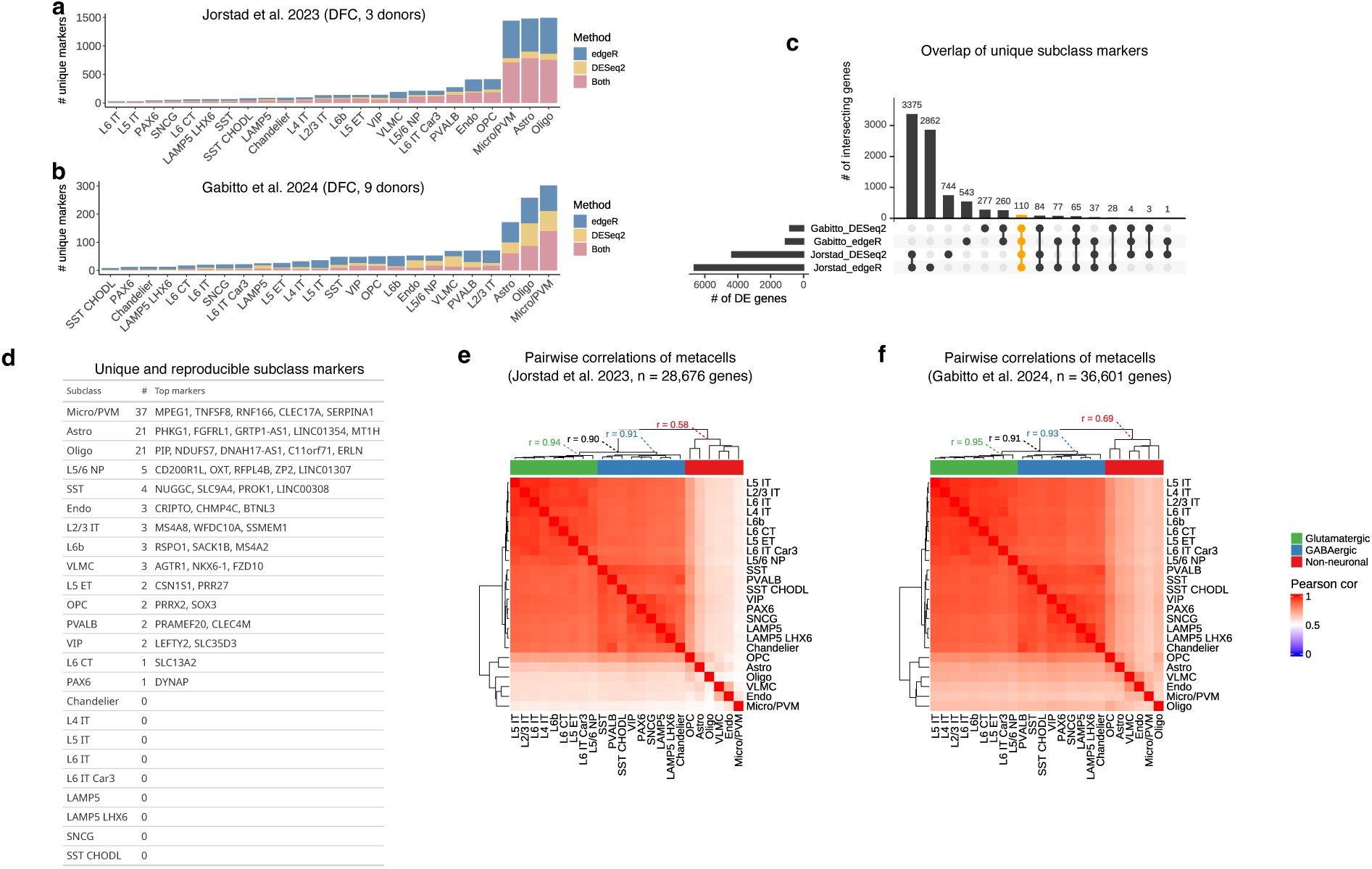
Cell type marker genes in human frontal cortex vary by choice of study and algorithm. **a-b)** Number of unique marker genes (FDR < .05) identified by differential expression (DE) analysis of snRNA-seq data from two studies^29,30^ of normal adult human dorsal frontal cortex (DFC) using `edgeR`^33^, `DESeq2`^34^, or both. **c)** Overlap of unique marker genes from **a** & **b** for all cell types. **d**) Unique and reproducible marker genes for each cell type ranked by significance. **e-f**) Heatmaps of genome-wide correlations among pseudobulked cell types from Jorstad et al.^30^ (**e**) and Gabitto et al^29^. (**f**). Mean correlations among metacells of a given class (glutamatergic, GABAergic, neuronal, non-neuronal) are reported at dendrogram branch points.

### Genome-wide expression levels are poorly explained by cell-type abundance in pseudobulk datasets

To further analyze the contributions of individual cell types to the expression levels of individual genes, we created pseudobulk samples using snRNA-seq data from individual donors and cortical regions^30^. To create each pseudobulk sample, we randomly sampled nuclei from all subclasses, recorded their identities, and summed the expression levels for each gene over all sampled nuclei. Using large pseudobulk datasets (n=1,518 samples), we then performed multiple linear regression to model gene expression as a function of subclass abundance, with each gene serving as the response variable, subclass abundance vectors (n=24) as predictors, and adjusted R^2^ and root mean-squared error (RMSE) as model outputs (**Fig. 2a**). We found that variation in subclass abundance explained only 7.6–8.2% of genome-wide expression variation (n=17,753 genes) in DFC samples from three donors (**Fig. 2b, Table S6**). To determine if more granular cell type definitions would produce better results, we constructed pseudobulk datasets from the same snRNA-seq data by randomly sampling supertypes (n=136) instead of subclasses and repeated the analysis. We found that variation in supertype abundance explained only 8.8–9.6% of genome-wide expression variation (**Fig. 2b, Table S6**). We repeated these analyses for additional cortical regions from the same donors. In MTG, we found that 7.5–8.1% of genome-wide expression variation was explained by variation in subclass abundance and 8.5–9.5% by supertype abundance (**Fig. S3a, Table S6**). In primary visual cortex (V1), these numbers were even lower (5.1–6.6% for subclass and 6.7–8.2% for supertype; **Fig. S3c, Table S6**).

**Fig. 2.**
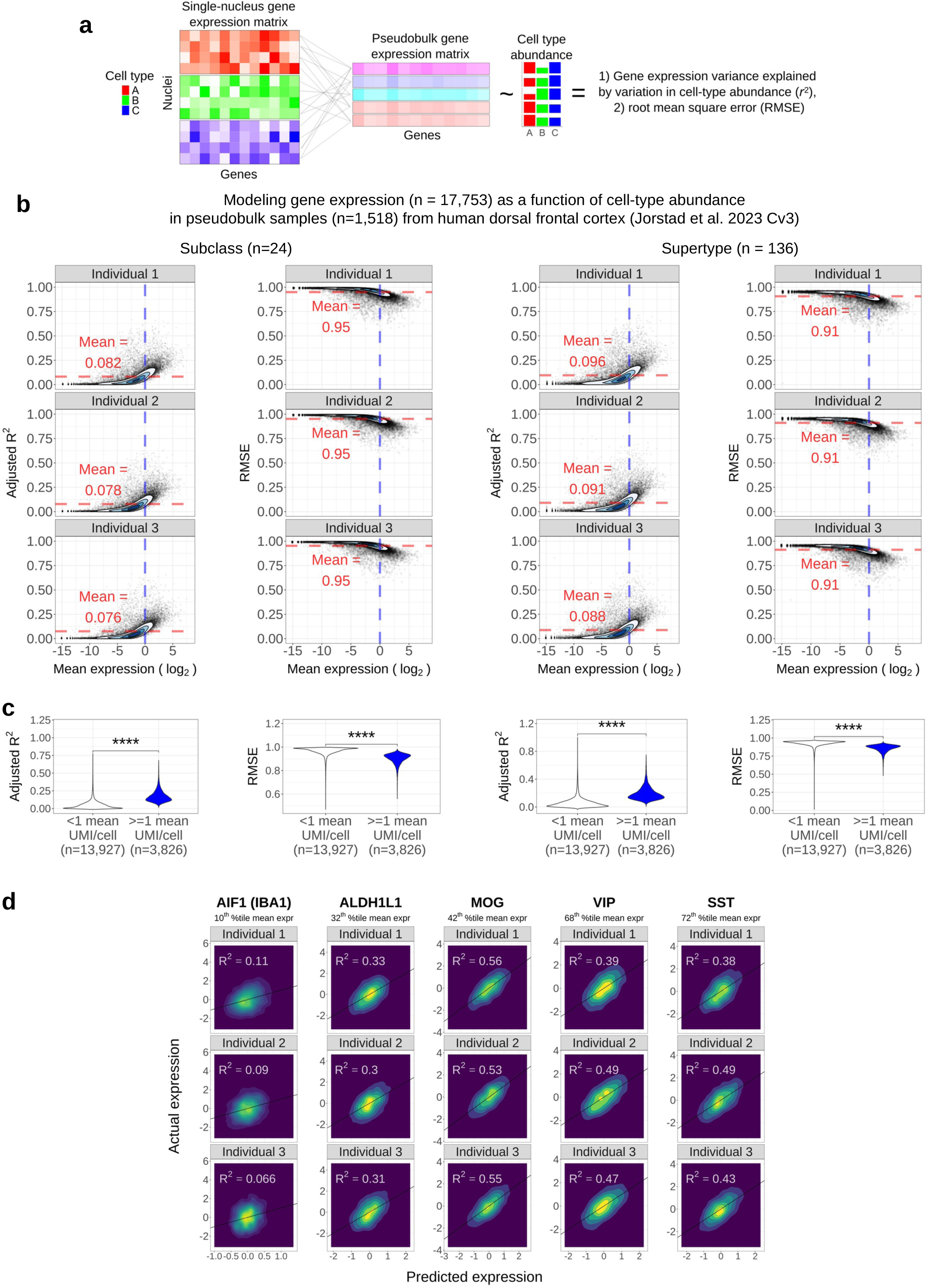
Genome-wide expression variation in pseudobulked snRNA-seq data from human frontal cortex is poorly explained by cell-type abundance, particularly for low-expressed genes. **a**) Schematic of pseudobulk modeling approach. Pseudobulk samples (n=1,518) were created by randomly sampling 10% of Cv3 DFC nuclei from each donor in Jorstad et al.^30^, recording their identities, and summing expression levels for each gene. Each gene was then used as the response variable in a multiple linear regression model with the abundance of each subclass (left) or supertype (right) as predictors. **b**) Adjusted R^2^ and root mean square error (RMSE) modeling results for all genes vs. mean expression level (log_2_). Red dotted lines denote mean modeling performance, while blue dotted lines denote mean expression of 1 UMI/nucleus^19,20^. **c**) Violin plots of modeling performance from (**b**), stratified by genes below and above this critical threshold. ****P < 0.0001. **d**) Predicted (fitted) vs. actual expression of select marker genes with different expression levels.

To validate these findings in a second snRNA-seq dataset^29^, we repeated pseudobulk modeling using DFC samples from nine additional donors and found that 6–10% of genome-wide expression variation was explained by variation in subclass abundance and 7.3–13% by supertype abundance (**Fig. S4, Table S7**). To determine whether deeper sequencing of nuclei would produce better results, we analyzed Smart-SEQ v4 (SSv4) snRNA-seq data^30^ from the same individuals and regions shown in **Fig. 2b** (DFC) and **Fig. S3a** (MTG). Despite >100x the read depth of Cv3 snRNA-seq data (**Table S1**), only 4.9–6.6% of genome-wide expression variation in DFC or MTG samples analyzed with SSv4 was explained by variation in subclass abundance and only 6.9%– 13% by supertype (**Fig. S5, Table S6**).

Previous studies have demonstrated that very low gene expression measurements in single cells (< 1 UMI / cell, on average) are generally unreliable for inferring meaningful cell-to-cell variation in expression levels^19,20^. In every pseudobulk analysis, we observed that modeling performance was significantly worse for genes expressed below this threshold, which represented 74.2–82.5% of all genes in Cv3 datasets and 18.8–28.9% of all genes in SSv4 datasets (**Figs. 2c, S3b&d, S4b, S5b&d)**. We further explored this finding at the level of individual subclass markers and observed better modeling performance for more highly expressed marker genes (**Fig. 2d**). However, even for the most deeply sequenced genes and datasets (SSv4), modeling performance was still on average quite poor. Collectively, these results indicate that human neocortical cellular identities are poor predictors for the expression levels of most genes in current snRNA-seq datasets.

### Covariation Projection Analysis (CoPA) reveals highly conserved modular gene activity in bulk and snRNA-seq datasets

The difficulty identifying reproducible marker genes for human neocortical subclasses may reflect the noisy nature of individual gene expression vectors in snRNA-seq data. However, genome-wide similarities among subclass metacells (**Fig. 1e-f, Fig. S2e-f**) strongly suggest that most gene activity in human neocortex is not cell-type-specific, which begs two important questions: How do shared genomic programs vary among neocortical cell types? And how are shared genomic programs impacted by disease? To address these questions, we searched for highly conserved gene coexpression modules in a large bulk RNA-seq dataset comprised of samples from multiple studies^38–44^ that surveyed genomic activity in neurotypical adult human DFC (**Table S8**). After preprocessing and batch correction^45,46^, the combined dataset consisted of 1,518 samples representing billions of cells, providing enormous statistical power to resolve highly reproducible gene coexpression modules in human neocortex.

We used an iterative version of a previously described module detection algorithm^22^ (**Fig. 3a**) to identify 1,016 fine-grained gene coexpression modules (**Table S9**), which were numbered in the order they were identified. Each module was summarized by its first principal component, or module eigengene (ME), and the similarity between each gene and each ME, or *k*_ME_, was quantified by Pearson correlation^47^. Following these steps, we produced a ‘snapshot’ for each module featuring several visualizations (**Fig. 3b-f**). First, we examined module coherence by calculating the percent variance explained by the ME and plotting z-scored expression patterns of the top 10 module genes (ranked by *k*_ME_) for 100 randomly selected samples from our combined dataset (**Fig. 3b**). Second, we assessed module reproducibility by plotting the distributions of pairwise correlations for module seed genes among samples from each of the input datasets we analyzed. These distributions were compared to empirical null distributions of pairwise correlations for randomly sampled genes (equivalent in number to module size) (**Fig. 3c)**. Third, we characterized module functions via enrichment analysis with a large collection of gene sets from the Molecular Signatures Database^48^ and OMICON (see companion study by Eliscu et al.) (**Fig. 3d**).

**Fig. 3.**
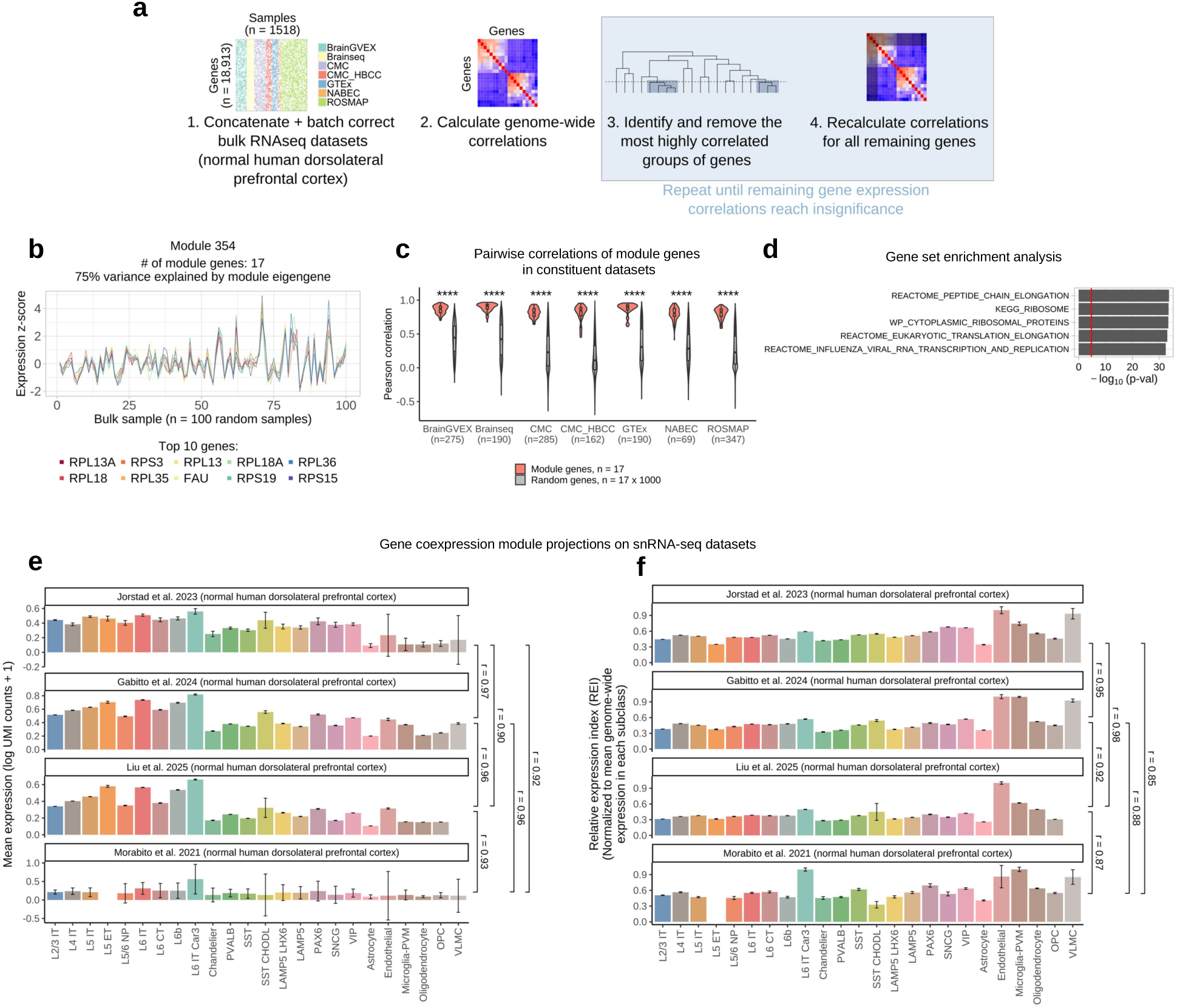
Covariation Projection Analysis (CoPA) reveals the cellular origins of bulk gene coexpression modules. **a**) Strategy for identifying gene coexpression modules in a large bulk RNAseq dataset comprising neurotypical adult human DFC samples (n=1,518) from six studies^39–44^. **b-f)** ‘Snapshot’ of one bulk gene coexpression module (Module 354). **b**) Expression patterns (z-scores) for module genes (n = 17) in 100 random DFC samples; ‘module eigengene’ is PC1 of the gene coexpression module^47^. **c**) Pairwise correlations of module genes in each of the input datasets referenced in (**a**) vs. pairwise correlations of random genes. **d**) Top results from enrichment analysis (one-sided Fisher’s exact test) of module genes using a large collection of gene sets from the Molecular Signatures Database^48^ and OMICON (https://theomicon.ucsf.edu). **e-f**) Module projections onto pseudobulked cell types from four snRNA-seq studies of human DFC^29–31,49^. Projections were calculated by averaging the expression of all module genes (natural log of UMI counts + 1) over all cells from each cell type in each dataset (**e**). A relative expression index (REI) was calculated by dividing values in (**e**) by the mean expression of all genes for each subclass, then scaling values to a maximum of 1 (**f**). Pearson correlations of projections are shown at right.

To elucidate the cellular origins of bulk coexpression modules, we projected module identities onto four snRNA-seq datasets from normal adult human DFC^29–31,49^. These datasets collectively represent 1,317,955 nuclei from 74 individuals sequenced to an average depth of 11,778 UMIs per nucleus (**Table S1**). All datasets but one used the BICCN reference taxonomy^32^ for cell type annotation. To harmonize annotations for the remaining dataset^49^, we assigned each nucleus to a BICCN subclass based on its maximum correlation with subclass metacells from Gabitto et al.^29^ (**Fig. S6**). To project bulk coexpression modules onto snRNA-seq subclasses, we first calculated the mean expression level (log-transformed UMI counts) for all module genes in all nuclei belonging to each subclass. This analysis revealed a common pattern for many modules, with highest expression in excitatory neurons, intermediate expression in inhibitory neurons, and lowest expression in non-neurons (**Fig. 3e**). Averaging genome-wide expression levels for each subclass confirmed this overall pattern (**Fig. S7**), which was consistent across cortical regions and reflects differences in the sizes of subclass nuclei and their overall transcriptional outputs. We therefore normalized each module projection by dividing the mean expression of module genes by the mean expression of all genes for each subclass and scaling all subclass values to lie between (0,1). This ‘relative expression index’ (REI) therefore quantifies the mean expression of module genes relative to the mean expression of all genes for each subclass (**Fig. 3f**). REI values for all modules and subclasses are provided in **Table S10**.

Snapshots of four ‘archetypal’ modules are shown in **Fig. 4a-p**, representing gene expression in all subclasses (column 1), all neuronal subclasses (column 2), a single subclass (column 3), or multiple non-neuronal subclasses (column 4). Although module coherence declined as expected with increasing module number (**Fig. 4a-d**), module reproducibility generally remained quite high (**Fig. 4e-h**). Many modules were significantly enriched with multiple gene sets (**Fig. 4i-l**) whose functions were consistent with module gene identities and projection patterns. We also observed that module projection patterns were very consistent for independent snRNA-seq datasets; in many cases, even subtle differences in REI values were extremely reproducible (**Fig. 4m-p**). Overall, 12,908 genes (68% of our input dataset) were identified as module seed genes (mean: ∼13 seed genes / module) (**Fig. 4q, Table S9**). Using an expanded module definition based on *k*_ME_ values (‘topmodposbc’), 17,972 genes (95% of our input dataset) were assigned to modules (mean: ∼18 genes / module) (**Fig. 4q, Table S9**). The mean pairwise correlation of module seed genes ranged from ∼0.95 for the first module to ∼0.37 for the last module (**Fig. 4r**). Over all modules, the mean correlation of REI projection patterns among all snRNA-seq datasets was 0.87 (**Fig. 4s**). All module snapshots can be browsed or searched on the CoPA Cabana web site that accompanies this study (https://oldhamlab.shinyapps.io/copacabana/), which also includes detailed descriptions of all gene sets used for enrichment analysis.

**Fig. 4.**
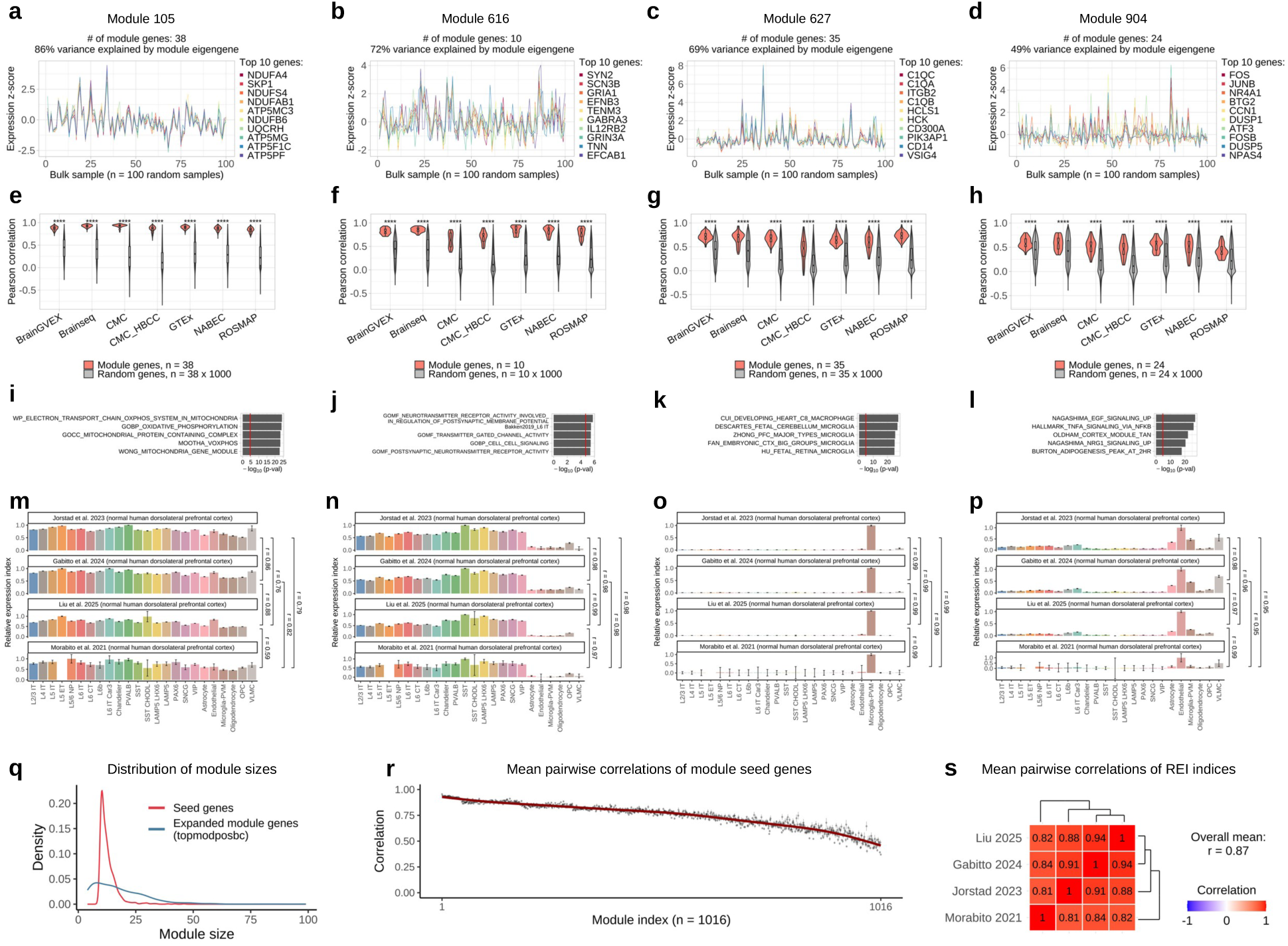
Bulk gene coexpression modules in human neocortex reflect highly reproducible patterns of genomic activity among neocortical cell types. **a-p)** Snapshots of four representative gene coexpression modules**. a-d**) Expression patterns (z-scores) for module genes in 100 random DFC samples. **e-h**) Pairwise correlations of module genes in each of the input datasets (Fig. 3a) vs. pairwise correlations of random genes. **i-l**) Top results from enrichment analysis (one-sided Fisher’s exact test) of module genes using a large collection of gene sets from the Molecular Signatures Database^48^ and OMICON (https://theomicon.ucsf.edu). **m-p**) Module projections onto pseudobulked cell types from four snRNA-seq studies of human DFC^29–31,49^, with relative expression indices calculated as described in Fig. 3. **q**) Sizes for all modules (n=1,016) using two module definitions (seed: initial set of module genes identified by the algorithm in Fig. 3a and used to form the module eigengene; topmodposbc: all genes that were positively and maximally correlated with the module eigengene below a Bonferroni-corrected significance threshold). **r**) Median pairwise correlations of seed genes for all modules. Vertical bars: +/- two S.E.; red line: smoothing spline. **s**) Mean pairwise correlations of projection indices (REI) from the four snRNA-seq datasets featured in (**m-p**) for all modules.

To systematically categorize module projection patterns and visualize the overall organization of gene activity in human neocortical subclasses, we performed consensus clustering of CoPA projection patterns for DFC snRNA-seq datasets from Jorstad et al.^30^ and Gabitto et al.^29^ (**Fig. 5a**). Specifically, by clustering modules using 1 – the minimum Pearson correlation of REI projections from both datasets, we identified 100 ‘meta-modules’ representing robust patterns of gene activity in human DFC (**Fig. 5b, Table S10**). Each meta-module represented on average ∼1% of all modules and ∼1% of all module genes (**Fig. 5b**). To visualize the expression patterns of meta-modules, we formed ‘eigenmodules’ by calculating PC1 of the REI projection patterns for merged modules in each snRNA-seq dataset. Visualizing these eigenmodules as heat maps revealed highly similar patterns of gene activity for meta-modules in both DFC snRNA-seq datasets (**Fig. 5c,d**). We also performed consensus clustering of cell types using CoPA projection patterns (i.e., the transpose of module clustering) (**Fig. 5a**), which recapitulated known relationships among subclasses (**Fig. 5c,d**). A parallel analysis of two MTG snRNA-seq datasets^29,30^ produced strikingly similar results (**Fig. S8, Table S10**).

**Fig. 5.**
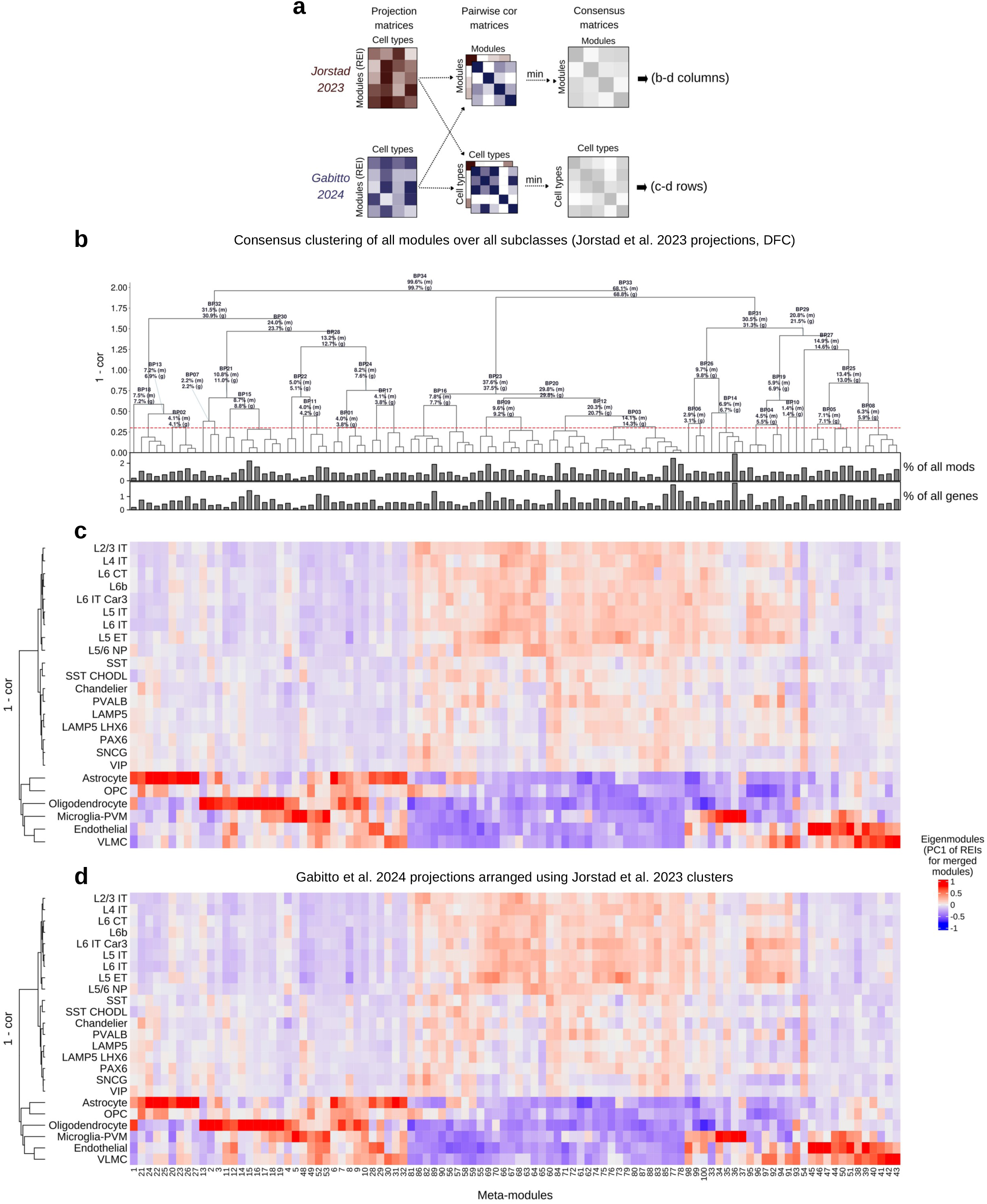
Clustering CoPA projection patterns reveals the major motifs of gene activity in human dorsal frontal cortex (DFC) cell types. **a**) Schematic of CoPA projection clustering. Left: projection matrices (REI vectors for 1,016 modules and 24 cell types) from two snRNA-seq datasets^29,30^. Middle: pairwise correlation (cor) matrices for modules (top) and cell types (bottom) derived from projection matrices. Right: consensus matrices for modules (top) and cell types (bottom) formed by retaining the minimum cor from both datasets at each position. Consensus cell types and modules were clustered using 1-cor and complete linkage. Meta-modules were identified by cutting the dendrogram at a height equal to the top 5% of pairwise correlations to merge modules (minimum meta-module size: three modules). **b**) Hierarchical clustering of meta-modules (n=100) using DFC snRNA-seq data from Jorstad et al^30^. Barplots below the dendrogram report the percentages of all modules (top) and genes (bottom) comprising each meta-module. Branch points above the red dashed line report the percentages of all constituent modules (‘m’) and genes (‘g’). **c-d**) Heatmaps of eigenmodules (first principal components of REI vectors for merged modules) produced using DFC snRNA-seq data from Jorstad et al.^30^ (**c**) and Gabitto et al^29^. (**d**). Cell types were clustered as described in (**a**).

By calculating the number of modules and genes associated with each meta-module and merged meta-modules (barplots and branch labels in **Fig. 5b**), we quantified the frequencies of expression patterns associated with cellular identities at different resolutions in human DFC (**Fig. 5b-d**). The initial bifurcation in meta-module clustering revealed that approximately half of all meta-modules have higher relative expression in neurons and half in non-neurons. On average, among non-neurons, vascular cells (endothelial + VLMC) had the highest relative expression for 17.5% of all meta-modules, oligodendrocytes for 16%, astrocytes for 15%, microglia for 11.5%, and OPCs for 2.5%. Among excitatory neurons, L5 ET neurons had the highest relative expression for 11% of all meta-modules, followed by 4% for L6 IT Car3, 3.5% for L2/3 IT, 3.5% for L5/6 NP, 2.5% for L4 IT, 1% for L5 IT, 1% for L6 IT, 0.5% for L6b, and 0% for L6 CT. Among inhibitory neurons, PVALB neurons had the highest relative expression for 3.5% of all meta-modules, followed by 2.5% for SNCG, 1.5% for SST, 1% for Chandelier, 1% for LAMP5, 0.5% for LAMP5 LHX6, 0.5% for SST CHODL, and 0% for PAX6 and VIP. Analysis of MTG produced very similar results (**Fig. S8**).

### dCoPA reveals coordinated and reproducible dysregulation of modular gene activity in neuropathologies

The consistency of modular gene expression patterns among neocortical subclasses (**Fig. 4s**) provides a novel framework for identifying transcriptional perturbations associated with disease. Specifically, we reasoned that comparing CoPA projection patterns for aggregated subclass nuclei derived from normal and pathological human brain samples could reveal cell-type-specific dysregulation of modular gene activity. By comparing the expression levels of coexpressed genes in all nuclei from each subclass, this strategy retains the statistical power of bulk coexpression analysis while mitigating the noise and sparsity inherent to snRNA-seq data and preserving cell-type-specific information. We therefore developed an empirical approach to identify bulk coexpression modules whose expression levels differed significantly in one or more neocortical subclasses between normal and pathological samples. We imposed two criteria: i) the Euclidean distance between the mean expression levels of module genes in case vs. control subclass nuclei must be greater than expected by chance, and ii) all genes in the module must change expression in the same direction (**Fig. 6a**).

**Fig. 6.**
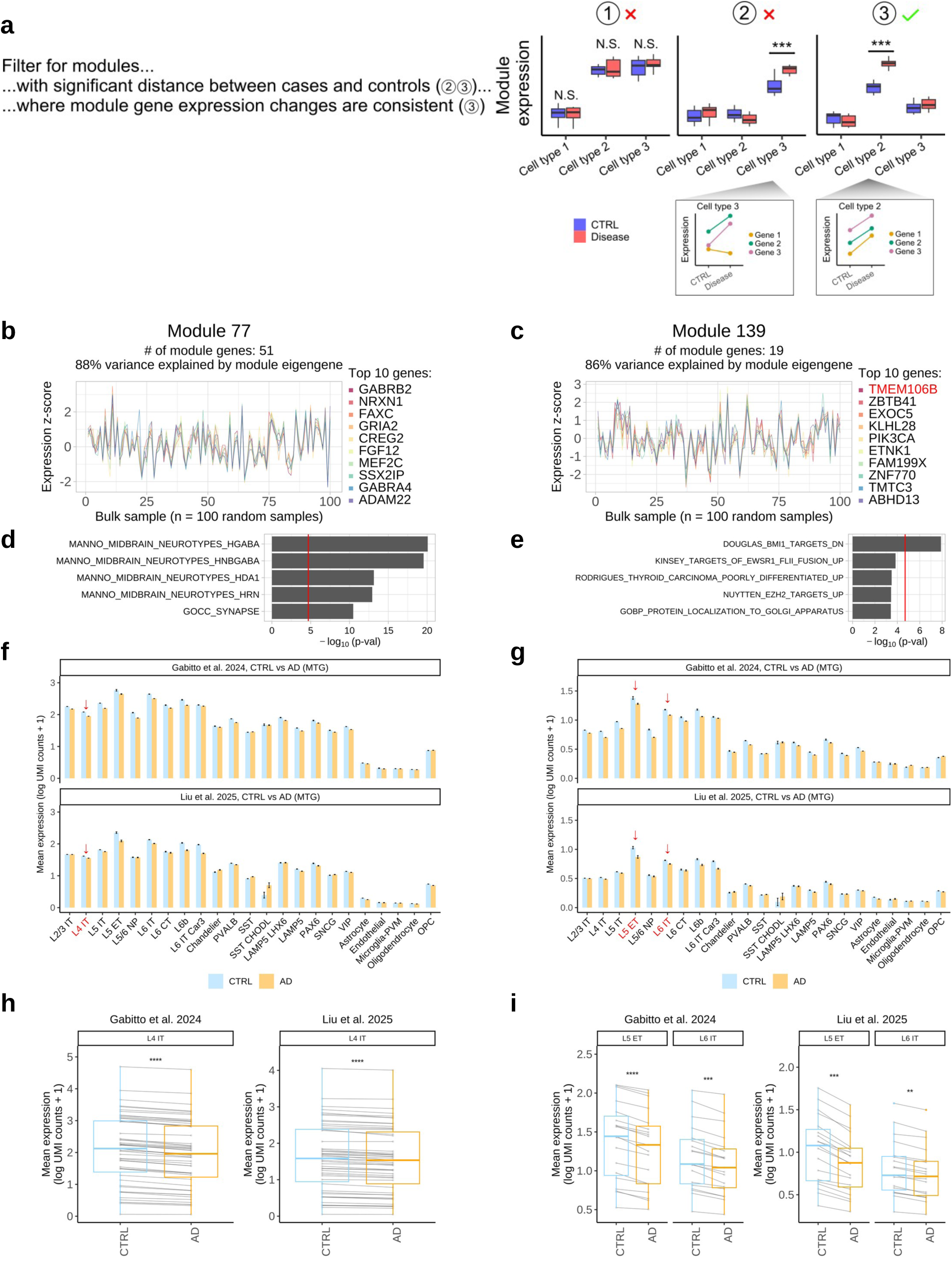
Differential CoPA (dCoPA) identifies gene coexpression modules that are significantly and uniformly dysregulated in specific cell types in disease. **a**) Schematic of dCoPA module identification strategy, which uses two criteria: i) the Euclidean distance between the mean expression levels of all module genes (averaged over all subclass nuclei) in cases vs. controls (CTRL) must be greater than expected by chance, and ii) all module genes must change expression in the same direction. **b-i**) Examples of two modules that meet dCoPA filtering criteria for Alzheimer’s disease (AD). **b-c**) Expression patterns (z-scores) for module genes in 100 random bulk DFC samples from our input dataset (Fig. 3a). **d-e**) Top results from enrichment analysis (one-sided Fisher’s exact test) of module genes using a large collection of gene sets from the Molecular Signatures Database^48^ and OMICON (https://theomicon.ucsf.edu). **f-g**) Projections of module genes onto two snRNA-seq datasets that analyzed CTRL and AD patient samples^29,49^, split by diagnosis. Significant cell types are marked by red arrows. **h-i**) Mean expression levels of all module genes in AD and CTRL nuclei for each cell type marked in (**f-g**). Asterisks indicate the empirical significance of module expression differences between AD and CTRL (compared to 10,000 randomly generated modules of the same size). *P < 0.05, **P < 0.01, ***P < 0.001, ****P < 0.0001.

We evaluated our approach using two large snRNA-seq datasets^29,49^ comprising DFC and MTG nuclei from neurotypical adults (CTRL) and individuals with Alzheimer’s disease (AD). Combined, these studies profiled 629,013 DFC nuclei / 318,994 MTG nuclei from 64 CTRL subjects and 1,478,438 DFC nuclei / 1,273,567 MTG nuclei from 126 AD subjects (**Table S1b**). We performed dCoPA separately for each study and brain region using real modules (n=1,016; **Fig. 4**) or random modules (i.e., modules formed by randomly selecting genes to match real module sizes). Importantly, after correcting for multiple comparisons, no random modules were significant in any analysis (**Fig. S9**). In contrast, over a third of real modules were significant for at least one cell type in each analysis, with a mean of about two significant cell types per significant module (**Fig. S9**).

To ensure the robustness of our findings, we stipulated that significant modules must also be reproduced in both snRNA-seq datasets. More specifically, for each brain region, we compared dCoPA results and retained only those significant modules for which all genes moved in the same direction in the same cell type in both snRNA-seq datasets^29,49^. Snapshots of two such modules are shown in **Fig. 6b-i**. Module 77 was significantly and reproducibly downregulated in L4 IT excitatory neurons from AD samples (**Fig. 6f,h**), while module 139 was significantly and reproducibly downregulated in L5 ET and L6 IT excitatory neurons from AD samples (**Fig. 6g,i**). We note that the top gene in this module (*TMEM106B*, **Fig. 6c**) has been reproducibly implicated in AD pathology via multiple genetic association studies^50–52^. Importantly, although all 70 genes in both modules were consistently and reproducibly downregulated in the same neuronal subclasses, none of these genes were significant by DE analysis of either snRNA-seq dataset using either method (**Table S11**).

To summarize and compare dCoPA results between snRNA-seq datasets (Gabitto et al.^29^ and Liu et al.^49^) and brain regions (MTG and DFC), we visualized the number of significant modules for each cell type as dot plots and separated modules that were significantly downregulated in AD nuclei from those that were significantly upregulated (**Fig. 7a,b** & **Fig. S10a,b**). For example, ‘CTRL modules | Gabitto SN’ (**Fig. 7a**) depicts the number of bulk coexpression modules from our integrated control cohort that were significantly downregulated in AD MTG nuclei vs. CTRL MTG nuclei for each cell type using snRNA-seq data from Gabitto et al. Interestingly, the overwhelming majority of significant modules identified by dCoPA were downregulated in AD nuclei (primarily in neurons), while the smaller number of significant modules that were upregulated in AD nuclei were more likely to occur in non-neurons (**Fig. 7a,b** & **Fig. S10a,b**).

**Fig. 7.**
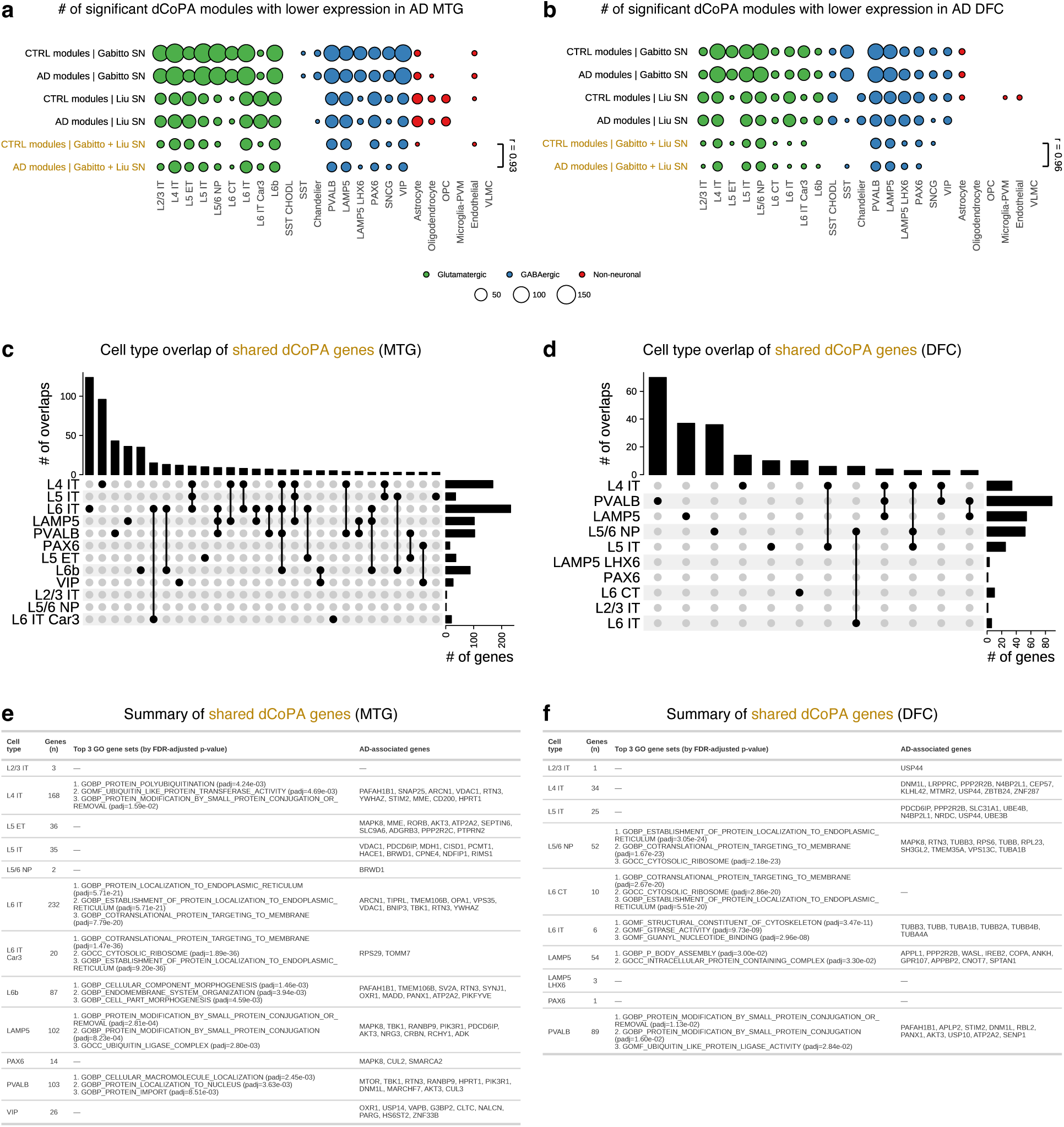
dCoPA reveals reproducible downregulation of modular gene activity in deep-layer intratelencephalic excitatory neurons, PVALB inhibitory neurons, and LAMP5 inhibitory neurons in Alzheimer’s disease (AD). **a-b**) Dotplots depict the number of bulk coexpression modules identified by dCoPA that were significantly down-regulated in AD vs. CTRL cell types from MTG (**a**) or DFC (**b**) in snRNA-seq data from Gabitto et al.^29^, Liu et al.^49^, or both (gold rows). CTRL / AD modules denote bulk coexpression modules derived from CTRL or AD samples, respectively. **c-d**) Cell type overlap of shared dCoPA genes from (**a**) and (**b**) (i.e., the intersection of all genes comprising the modules in the bottom two rows in each figure panel). **e-f**) Top three results from Gene Ontology^54^ enrichment analysis for genes featured in (**c-d**) are shown in third column, and AI-powered multi-agent search results for connections to AD biology are listed in the fourth column (see also **Table S14**). Interactive dot plots (**a-b**) are available on the CoPA Cabana web site (https://oldhamlab.shinyapps.io/copacabana/).

In principle, the bulk coexpression modules used as input for dCoPA can be derived from normal or pathological cohorts. We therefore reran CoPA to identify bulk coexpression modules in the ROSMAP cohort of AD patient samples (n = 248 from DFC)^53^. This analysis identified 1,010 modules (**Table S12**), which were projected onto AD and CTRL nuclei from Gabitto et al.^29^ or Liu et al.^49^ in the same fashion as the bulk CTRL modules (**Fig. 7a,b** & **Fig. S10a,b**). We next compared the number of bulk CTRL or bulk AD modules that were significantly and reproducibly downregulated in each cell type (bottom two rows in **Fig. 7a,b**). Even though the bulk CTRL and bulk AD modules were independently derived, the overall pattern of cell-type-specific downregulation identified by dCoPA was extremely reproducible (**Fig. 7a,b**: r = 0.93 for MTG and r = 0.96 for DFC), with the greatest number of downregulated modules observed in deep layer intratelencephalic excitatory neurons and PVALB or LAMP5 inhibitory neurons from both regions. dCoPA results for all modules and comparisons are provided in **Table S13**, while **Table S14** lists all genes comprising the significant and reproducible modules identified by dCoPA. Users can also interact with the dot plots in **Fig. 7a,b** & **Fig. S10a,b** on the CoPA Cabana web site (https://oldhamlab.shinyapps.io/copacabana/), where clicking on a dot reveals the snapshots of modules that are significantly and reproducibly dysregulated in a given cell type.

We next compared the identities of genes comprising the bulk CTRL and bulk AD modules that were significantly and reproducibly downregulated in each cell type. We observed the most significant overlap (P < 1.0E-14) of these genes in L4 IT, L5 ET, L5 IT, L6 IT, L6 IT Car3, L6b, PVALB, and LAMP5 neurons, with many of the same genes downregulated in multiple neuronal cell types (**Fig. S11**). To identify the most reproducible set of affected genes, we took the intersection of all genes comprising bulk CTRL or bulk AD modules that were significantly and reproducibly downregulated in each cell type (bottom two rows in **Fig. 7a,b**). We then evaluated how these genes were distributed among cell types and identified deep layer intratelencephalic excitatory neurons and PVALB or LAMP5 inhibitory neurons as the most affected, albeit with some variability between brain regions (**Fig. 7c,d**). Enrichment analysis revealed significant over-representation of Gene Ontology^54^ categories related to ribosomal function, protein localization, protein conjugation, and protein ubiquitination among module genes that were coordinately and reproducibly downregulated in specific neuronal subtypes (**Fig. 7e,f**: left). We also performed an AI-powered multi-agent search of the biomedical literature to identify studies that have implicated these genes in AD biology (**Fig. 7e,f**: right; references for these studies are provided in **Table S15**). To evaluate dCoPA for a different type of neuropathology, we obtained additional bulk and snRNA-seq data from patients with schizophrenia (SCZ) and controls. We analyzed the same set of bulk CTRL modules as we did for the AD analysis (n = 1,016), while bulk SCZ modules (n = 1,083) were identified by applying CoPA to 171 DFC samples from SCZ patients from the Brainseq Phase 1 study^43^ (**Table S8** and **Table S16**). Using dCoPA, we analyzed the CTRL and SCZ modules in two snRNA-seq datasets that collectively profiled 461,353 nuclei from 77 CTRL subjects and 385,547 nuclei from 71 SCZ subjects^55^ (**Table S1**). As with AD, the vast majority of significant and reproducible modules identified by dCoPA were downregulated in SCZ neurons, while the few that were upregulated in SCZ were primarily observed in non-neurons (**Fig. 8a,b**). We then compared the number of bulk CTRL or bulk SCZ modules that were significantly and reproducibly downregulated in each cell type and again found that the overall pattern of cell-type-specific downregulation identified by dCoPA was highly reproducible (r = 0.78; bottom two rows in **Fig. 8a,b**).

**Fig. 8.**
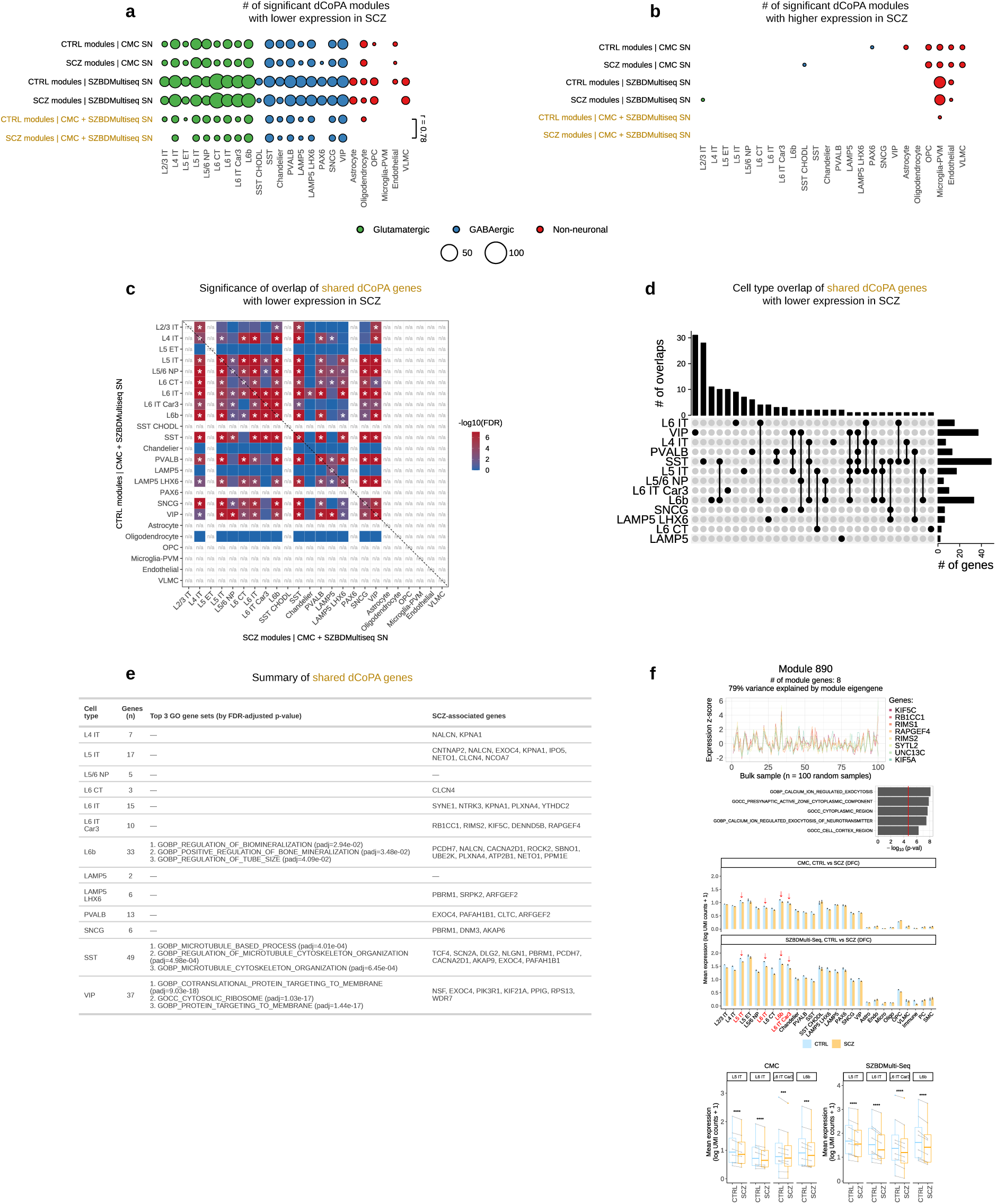
dCoPA reveals reproducible downregulation of modular gene activity in SST inhibitory neurons, VIP inhibitory neurons, and L6b excitatory neurons in schizophrenia (SCZ). **a-b**) Dotplots depict the number of bulk coexpression modules identified by dCoPA that were significantly down-regulated (**a**) or up-regulated (**b**) in SCZ vs. CTRL cell types from DFC in snRNA-seq data from the SZBDmulti-seq cohort^65^, the CMC cohort^42^, or both (gold rows). CTRL / SCZ modules denote bulk coexpression modules derived from CTRL or SCZ samples, respectively. **c)** Significance of overlap (one-sided Fisher’s exact test) for shared dCoPA genes with lower expression in SCZ (i.e., the intersection of all genes for each subclass in the bottom two rows in [**a**]). **d)** Cell type overlap of shared dCoPA genes with lower expression in SCZ (i.e., the same genes analyzed in [**c**]). **e**) Top three results from Gene Ontology^54^ enrichment analysis for genes featured in (**c-d**) are shown in third column, and AI-powered multi-agent search results for connections to SCZ biology are listed in the fourth column (see also **Table S14**). **f**) Example of a module exhibiting uniform and reproducible downregulation in deep-layer excitatory neurons in SCZ. Interactive dot plots (**a-b**) are available on the CoPA Cabana web site (https://oldhamlab.shinyapps.io/copacabana/).

We next compared the identities of genes comprising the bulk CTRL and bulk SCZ modules that were significantly and reproducibly downregulated in each cell type. We observed the most significant overlap (P < 1.0E-7) of these genes in L5 IT, L6 IT, L6 IT Car3, L6b, SST, and VIP neurons, with many of the same genes downregulated in multiple neuronal cell types (**Fig. 8c**). To identify the most reproducible set of affected genes, we took the intersection of all genes comprising bulk CTRL or bulk SCZ modules that were significantly and reproducibly downregulated in each cell type (bottom two rows in **Fig. 8a**). We then evaluated how these genes were distributed among cell types and identified SST, VIP, and L6b neurons as the most affected (**Fig. 8d**). Enrichment analysis revealed significant over-representation of Gene Ontology biological processes related to protein targeting to membrane and microtubules among module genes that were coordinately and reproducibly downregulated in these neuronal subtypes (**Fig. 8e** [left]). We also performed an AI-powered multi-agent search of the biomedical literature to identify studies that have implicated these genes in SCZ biology (**Fig. 8e**: right; references for these studies are provided in **Table S15**). An example of one module with significant and reproducible downregulation in multiple SCZ excitatory neuron subtypes is shown in **Fig. 8f**, with similar visualizations for all significant modules available in CoPA Cabana (https://oldhamlab.shinyapps.io/copacabana/). Interestingly, we noticed that some modules identified by dCoPA were significant for SCZ and AD. Specifically, we identified 18 examples of modules that were significantly, coordinately, and reproducibly downregulated in the same cell type(s) in AD and SCZ. This overlap was highly significant (P = 3.4e-24).

## DISCUSSION

CoPA is a novel and statistically motivated approach for combining the power of bulk tissue sampling with the precision of single-cell analysis to reveal the origins of highly reproducible gene coexpression modules and their relative importance among cell types. By applying CoPA to vast amounts of public gene expression data, we have shown that very few gene coexpression modules in human neocortex are cell-type-specific; however, subtle differences in modular gene activity among cell types are nevertheless extremely reproducible. This reproducibility enables comparisons of module projection patterns between single-cell datasets derived from normal and pathological human tissue samples with dCoPA. Using this approach, we identified gene coexpression modules that are uniformly and reproducibly dysregulated in specific neocortical cell types from patients with AD or SCZ. To share our findings, we have created a user-friendly R Shiny app called CoPA Cabana (https://oldhamlab.shinyapps.io/copacabana/) that can also perform CoPA for thousands of gene coexpression networks in OMICON (https://theomicon.ucsf.edu), which are derived from standardized analysis of 116 datasets representing >17K normal and neoplastic human brain samples (see companion study by Eliscu et al.).

Our central insight is that the population structure of genomic activity – efficiently discovered through covariation analysis of intact tissue samples – provides a natural framework for interpreting single-cell datasets. Because large bulk datasets may represent thousands of individuals and billions of cells, they provide enormous statistical power to discern highly reproducible modules of genomic activity, which also reduce the dimensionality of gene expression data. By projecting these modules onto pseudobulked cell types from single-cell datasets, CoPA clarifies the relative importance of these modules for each cell type, and also how cell types differ from one another. Given that neocortical subclass relationships were faithfully reconstructed by clustering bulk coexpression module projection patterns (e.g., **Fig. 5c**), it follows that bulk coexpression modules could potentially be used to refine cell-type annotations in neocortical single-cell datasets. However, any such effort should acknowledge that similarities far outweigh differences in gene expression among neuronal cell types in human neocortex. For example, the average genome-wide correlation among all neuronal subclasses (excitatory and inhibitory) in human DFC and MTG is ∼0.9; for oligodendrocytes and OPCs, the corresponding value is ∼0.7. Therefore, claims of ever more granular neuronal subtypes should be treated with caution unless they have been unambiguously replicated in multiple, independent datasets.

dCoPA provides a statistical framework for comparing CoPA projection patterns in single-cell datasets from different cohorts. Unlike standard DE analysis, which treats genes separately and ignores their coexpression relationships, dCoPA uses the covariation structure of genomic activity in bulk datasets as a scaffold for inquiry. By comparing the mean expression levels of coexpressed genes between vast numbers of cells or nuclei for each cell type and requiring that expression differences between cohorts are significant, uniform, and reproducible, we introduce a novel and robust strategy for revealing pathological gene activity in single-cell datasets. Application of this strategy to AD and SCZ produced several interesting findings.

First, in both AD and SCZ, dCoPA identified many more modules that were significantly downregulated than upregulated. Second, most of the downregulated modules implicated neurons, while most of the upregulated modules implicated non-neurons. Third, many significant modules were coordinately downregulated in multiple neuronal subclasses (often including excitatory and inhibitory subclasses). Fourth, there were more significant dCoPA modules in AD MTG than AD DFC, consistent with known AD pathology^56^. Fifth, implicated cell types were highly consistent whether bulk coexpression modules were derived from normal or pathological cohorts. And sixth, although the most affected cell types differed by pathology (**AD:** L6 IT, L4 IT, PVALB, LAMP5; **SCZ:** SST, VIP, L6b), a significant number of dCoPA modules were dysregulated in the same direction and the same cell type(s) in AD and SCZ.

Collectively, our findings highlight distinct gene coexpression modules that are selectively vulnerable to dysregulation in AD and SCZ. Hardly any of these modules are cell-type-specific, and most of the genes comprising them would not be identified by standard DE analysis of individual cell types (**Table S11**). However, these genes paint a detailed portrait of the cellular activities most impacted by disease. For example, the genes that are most recurrently identified by dCoPA for AD (**Table S14**) suggest a coordinated neuronal collapse of protein synthesis, membrane trafficking / lysosomal clearance, axonal / synaptic delivery, and nuclear gene regulatory control. In SCZ, the same analysis (**Table S14**) points to reduced expression of genes involved in synaptic structure, vesicle cycling, receptor localization, and dendritic spine structure. Our results also raise the tantalizing possibility that targeted reversal of dysregulated modules identified by dCoPA may restore normal cellular functions and ameliorate pathological symptoms. Our study has several important limitations. First, CoPA projections can be misleading if bulk coexpression modules are driven by cell types or states that are not represented in single-cell datasets. For example, none of the snRNA-seq datasets we analyzed included ependymal or choroid plexus cells. Second, the accuracy of module projections may be limited by the sample size available for each annotated cell type; in some snRNA-seq datasets, certain cell types were represented by < 200 nuclei (e.g., VLMC and SST CHODL), limiting power. Third, there are many ways to define gene coexpression modules in bulk datasets. We took a conservative approach and iteratively removed all groups of highly correlated genes without merging them, which means that some modules may be closely related and functionally redundant. However, dCoPA analysis of modules derived independently from normal or pathological bulk datasets revealed remarkable consistency in the patterns of affected cell types, underscoring the robustness of our approach.

In summary, we have described a novel analytical strategy that combines the power of bulk coexpression analysis with the precision of single-cell methods to reveal the cellular origins of highly reproducible modules of genomic activity in health and disease. We have also created the CoPA Cabana web site (https://oldhamlab.shinyapps.io/copacabana/), which accompanies this study and provides a novel resource for the neuroscientific community by hosting all CoPA and dCoPA results from our analyses. We expect that application of CoPA and dCoPA to other pathological conditions and omics dataset types will clarify the organization of genomic activity in disparate tissues and the combinations of molecular programs and cell types that are selectively impacted by disease.

## METHODS

All analyses were performed in R (v4.4.1) (https://cran.r-project.org/) or Python (v3.9.16).

### 1. snRNA-seq DE analysis

We downloaded two snRNA-seq datasets^29,30^ comprising ∼527K neocortical nuclei from dorsolateral prefrontal cortex (DFC) and middle temporal gyrus (MTG) of 12 neurotypical adult human donors (**Table S1**). Data were produced using the 10x Chromium V3 (Cv3) platform.

#### 1.1 Construction of donor- and subclass-level pseudobulk samples from snRNA-seq data

To perform differential expression (DE) analysis, we used a pseudobulk approach to avoid pseudoreplication bias and excessive false discoveries that can result from performing DE directly on individual cells or nuclei from the same donors^16–18^. Raw UMI counts were summed for each subclass (n = 24) from each donor (n = 12) in each brain region, resulting in 72 DFC or MTG pseudobulk samples from Jorstad et al.^30^ and 216 DFC or MTG pseudobulk samples from Gabitto et al^29^.

#### 1.2 Differential expression (one subclass vs. all other subclasses)

DE was performed using the R packages `edgeR`^33^ (v4.4.2) and `DESeq2`^34^ (1.46.0). For `edgeR` we used the glmLRT (Generalized Linear Model Likelihood Ratio Test) pipeline without `filterByExpr` and with `robust = TRUE` for the `estimateDisp` function. For DESeq2, `test = “LRT”` was utilized with the `reduced` model defined using donors only. DE was performed after filtering genes to those present in both Jorstad et al.^30^ and Gabitto et al.^29^ (n = 20,568 for DFC and n = 20,892 for MTG). For each subclass, marker genes were defined as those that were significantly upregulated (FDR < .05 and log_2_(fold change) > 0) and unique (i.e., significantly upregulated in only one subclass).

### 2. Pseudobulk modeling

In addition to the datasets used for DE analysis, we downloaded additional snRNA-seq data from neurotypical adult human neocortex sample cohorts from Jorstad et al.^30^, including Cv3 primary visual cortex (V1) (n = three donors and 141,836 nuclei), SMART-seq V4 (SSv4) DFC (n = three donors and 4,551 nuclei), and SSv4 MTG (n = three donors and 17,307 nuclei) (**Table S1**).

#### 2.1 Pseudobulk dataset creation for modeling gene expression vs. cell type abundance

For each snRNA-seq dataset cohort, pseudobulk samples were created by randomly sampling and summing 10% of all nuclei to create synthetic samples. We matched the number of samples (n = 1,518) and genes (n = 18,913) in each pseudobulk dataset to match the integrated bulk control dataset (**Fig. 3a**). The cell type composition of each pseudobulk sample was recorded to create cell type abundance vectors for linear modeling of gene expression.

#### 2.2 Linear modeling of pseudobulked gene expression by cell type abundance

To model pseudobulked gene expression as a function of cell type abundance, we performed multiple linear regression with each gene (0*_s_*) in each pseudobulk dataset consisting of *_s_* samples as response and cell type abundance vectors (*_a_*) as predictors using the lm function in R:

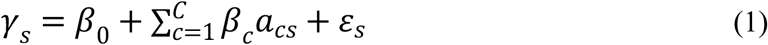

where *_acs_* represents the relative abundance of cell type *_c_* in sample *_s_*, *_βc_* represents the regression coefficient for cell type *_c_*, *_β_*_$_ represents the model intercept, and s*_s_* represents the error term. Cell types (*_c_*) were defined using either subclass (n = 24) or supertype (n = 136) definitions. Modeling performance was quantified with Adjusted R^2^ or the root mean squared error (RMSE):

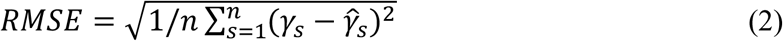

Where *γ_s_* represents real gene expression in sample *_s_*, *γ^_s_* represents predicted gene expression in sample *_s_*, and *n* represents the total number of samples.

### 3. CoPA

CoPA (**Co**variation **P**rojection **A**nalysis) is a pipeline for integrating genome-wide covariation analysis of bulk tissue samples with analysis of pseudobulked cell types from single-cell or single-nucleus datasets. The full pipeline starts with a large bulk dataset as input, followed by genome-wide covariation analysis, followed by covariation module projection onto pseudobulked cell types from single-cell or single-nucleus datasets (**Fig. 3a**).

#### 3.1 Bulk RNA-seq pre-processing

We downloaded bulk RNA-seq data from six studies^39–44^ representing adult (≥18 years) human neurotypical DFC samples. After filtering and quality control (see below), we obtained a total of 1,518 samples. All pre-processing was performed from FASTQ files with the exception of GTEx (see below). Raw sequence reads were downloaded from Synapse (https://www.synapse.org/) using the following identifiers: syn8612097 (Religious Orders Study and Memory and Aging Project Study, or ROSMAP), syn18134196 (Common Mind Consortium, or CMC, split between the MSSM/Penn/Pitt cohort and the NIMH Human Brain Collection Core, or HBCC, cohort), syn8227833 (BrainSeq Phase 1), and syn7062404 (BrainGVEX). Data from GTEx (v8) was downloaded from the AnVIL repository (https://anvil.terra.bio). Data from the North American Brain Expression Consortium (NABEC) was downloaded from dbGaP (https://dbgap.ncbi.nlm.nih.gov/) with the accession ID phs001353.v1.p1.

Reads were quantified from raw expression files using `kallisto`^57^ (v0.44.0) with default settings for paired-end data. A `kallisto` index was built using the GENCODE v39 transcriptome (GRCh38)^58^. BAM files from GTEx were converted to FASTQ files using the `run_SamToFastq.py` script from the Broad Institute’s `gtex-pipeline` repository (https://github.com/broadinstitute/gtex-pipeline). Transcript-level abundances were summarized to gene-level counts with `tximport`^59^ (v1.34.0) using a GENCODE-derived (v39) transcript-to-gene mapping.

Each bulk dataset was processed using the `SampleNetwork`^46^ R function, a bulk expression pre-processing tool designed for sample outlier removal and batch correction. Outliers were defined as samples with connectivity more than four standard deviations below the mean connectivity of all samples over all genes (*_Z_*. *_K_* < -4). Outliers were iteratively removed until no samples with *_Z_*. *_K_* < -4 remained. Next, samples were batch corrected for library batch if the corresponding information was available. Batch correction was performed using the `Combat`^45^ R function, which is built into `SampleNetwork`. Finally, genes with zero variance were removed. After outlier removal and batch correction, all datasets were concatenated into a single dataset after aligning to shared genes (n = 18,913). `SampleNetwork` was then applied again to the integrated dataset to remove outliers and to batch correct by individual dataset. The final batch-corrected, integrated bulk control dataset was used as input for downstream analyses.

In addition to the integrated bulk control dataset, two additional disease-specific bulk datasets were prepared and utilized for downstream analyses: 248 samples from ROSMAP^53^ with Alzheimer’s disease (AD) and 171 samples from Brainseq Phase 1^43^ with schizophrenia (SCZ). These datasets were quantified from FASTQ files using `kallisto` and processed by `SampleNetwork` in an identical fashion to each individual dataset of the integrated bulk control dataset.

#### 3.2 Unsupervised co-expression module detection with iterative sequestering

We used a variant of the coexpression module detection approach described by Kelley et al^22^. We implemented a ‘greedy’ approach that iteratively finds the most highly correlated groups of genes and removes them prior to the next iteration, thereby increasing the discoverability of modules that might be masked by more highly correlated groups of genes. Each iteration of the algorithm proceeded as follows: first, pairwise Pearson correlations were calculated for all genes over all samples of the integrated bulk control dataset (n = 1,518) to produce a genome-wide similarity matrix. Next, the similarity matrix was hierarchically clustered with complete linkage using the `flashClust`^60^ package (v1.1.2) using 1 − *_cor_* as the distance measure. The resulting dendrogram was then cut at a height corresponding to the top 0.01% of pairwise correlations to produce an initial set of modules, which were then filtered for modules with a minimum size of 10 genes - this threshold was enforced for all subsequent clustering steps of the algorithm. After recording this initial set of modules, the seed genes representing these modules were removed from the similarity matrix, thus concluding the iteration. The subsequent iteration would then proceed on the trimmed similarity matrix.

For a given dendrogram cut height, the number of modules found would naturally decrease with each iteration until no modules were found. At this point, the algorithm would relax the cut height parameter and then repeat the entire iterative loop again. Initial cut height was set at the top 0.01% of all pairwise correlations, then was lowered in a stepwise fashion according to the following thresholds: 0.1%, 1%; then increasing by 1% until 10%; then increasing by 5% each iteration until a statistically determined lower threshold for minimum module seed gene pairwise correlation was reached. This threshold was defined as the 99th percentile of the distribution of minimum pairwise correlations calculated from randomly generated modules of size 10 (n = 10,000). Once the cut height exceeded this threshold, the algorithm would stop, concluding the module discovery process.

Expanded module definitions (“topmodposbc”) were then calculated for all modules. First, each module was summarized by its module eigengene^47^, defined as the first principal component of the gene expression vectors defining the module. Genes were then ranked for each module by the WGCNA measure of intramodular connectivity (*_k_*_mE_, defined as the Pearson correlation of a gene with a module eigengene^47^), and kept if the *_k_*_mE_ value was significant according to a Bonferroni-corrected threshold (where the number of comparisons is defined by the number of genes times the number of modules). Finally, genes were kept for a given module if the gene had the highest *_k_*_mE_ value for that module compared to all others. Defining modules this way resulted in more genes associated with each module on average, but a fraction of the modules had fewer topmodposbc genes than the 10 gene threshold enforced for module seed genes (**Fig. 4q**).

Before continuing with downstream analyses, modules underwent additional quality control filtering. First, modules with three or fewer genes according to the topmodposbc expanded definition were discarded. This resulted in 135 modules discarded from the integrated bulk control modules, 117 modules discarded from the ROSMAP AD modules, and 64 modules discarded from the Brainseq SCZ modules. Second, for the integrated bulk control modules specifically, the significance of pairwise correlations of topmodposbc genes in each input dataset for the integrated bulk control dataset was calculated for all modules by comparing real pairwise correlations of a given module in a given dataset to the pairwise correlations of genes from randomly generated modules (n = 1,000), matched to the size of the module. Modules that did not have pairwise correlations significantly greater than chance in at least two datasets were discarded. This resulted in seven additional modules being discarded from the integrated bulk control modules. The final module count for each dataset was as follows: 1,016 modules for the integrated bulk control dataset, 1,010 modules for the ROSMAP AD dataset, and 1,083 modules for the Brainseq SCZ dataset.

#### 3.3 Gene set enrichment analysis

To clarify module identity and function, we performed enrichment analyses of module topmodposbc genes with two collections of gene sets: the Molecular Signatures Database (MSigDB v7.4)^48^ from the Broad Institute (https://www.gsea-msigdb.org/gsea/msigdb) and a curated list of cell type gene sets sourced from a wide range of bulk RNA expression and SC/SN RNAseq datasets available in OMICON (https://theomicon.ucsf.edu) (see companion study by Eliscu et al.). We used the following human collections from the MSigDB database: H (hallmark gene sets), C2 (curated gene sets), C5 (ontology gene sets) and C8 (cell type signature gene sets). Enrichment tests were performed with a one-sided Fisher’s exact test using the base R `phyper` function.

#### 3.4 Projecting modules onto snRNA-seq data

We projected co-expression modules onto a series of snRNA-seq datasets. In addition to the previously mentioned Jorstad et al.^30^ and Gabitto et al.^29^ datasets, we downloaded two additional datasets from studies of adult human neurotypical DFC. Data from Liu et al. 2025^49^ were downloaded from https://compbio.mit.edu/AD_Multiomic_MultiRegion/, consisting of 626,175 nuclei from 55 individuals. This resource aggregates nuclei generated by Liu et al. with previously published data from Mathys et al. (2023)^61^, Xiong et al.^62^, and Mathys et al. (2024)^63^. Data from Morabito et al. 2021^31^ were downloaded from the Gene Expression Omnibus (https://www.ncbi.nlm.nih.gov/geo/) using the accession GSE174367, consisting of 22,796 nuclei from seven individuals.

To project modules onto neocortical subclasses from snRNA-seq datasets, snRNA-seq data were log-transformed + 1 (log1p). Then, for each subclass and each module (defined using either seed genes or topmodposbc genes), mean expression was calculated over all nuclei from each subclass (**Fig. 3e**). To calculate the relative expression index (REI), these projection indices were then normalized by the mean expression of all genes in each subclass, then scaled so that the maximum REI value was 1 (**Fig. 3f**).

#### 3.5 Cross-dataset subclass mapping

To enable subclass-specific cross-dataset comparisons between Gabitto et al.^29^ and Liu et al.^49^, we mapped all nuclei from Liu et al. to Gabitto et al. subclasses using the following strategy: first, raw UMI counts for both datasets were log-transformed +1 (log1p). Then, metacells for each Gabitto et al. subclass (n = 23) were created by averaging expression vectors for all nuclei belonging to a given subclass to produce a gene by metacell matrix. Then, the Pearson correlations between each nucleus in Liu et al. and each metacell from Gabitto et al. were calculated, and subclass labels were mapped to each nucleus based on the most highly correlated metacell (**Fig. S6**). We repeated this process for DFC and MTG regions separately. VLMCs were not included in the final mapping due to the absence of that cell type in Liu et al.

#### 3.6 Consensus meta-module clustering

To identify patterns among module projections, we hierarchically clustered module projections to produce “meta-modules” that summarize common archetypes of gene activity across neocortical subclasses. Before doing so, to ensure that we identified projection patterns that were reproducible across datasets, we calculated a consensus minimum similarity matrix from REI projections of the integrated bulk control modules onto Jorstad et al.^30^ and control samples from Gabitto et al.^29^ (**Fig. 5a**). Specifically, for each dataset we calculated module-by-module similarity matrices by calculating pairwise Pearson correlations of all modules over all REI projection indices. The resulting two matrices were then collapsed into a single consensus matrix by calculating the parallel minimum across both matrices using the `pmin` R function, which in essence takes the minimum correlation value at each position after overlaying both matrices on top of one another. This consensus approach ensures that any co-expression relationships are reproducible across both input datasets.

This consensus similarity matrix was then processed using previously described methods. We clustered the consensus similarity matrix using the `flashclust` (v 1.1.2) implementation of complete linkage hierarchical clustering with 1 − *_cor_* as a distance measure. The resulting dendrogram was then cut at a height corresponding to the top 5% of all pairwise correlations, producing a series of meta-modules where each meta-module represents a group of highly correlated module projection patterns. Meta-modules were kept if the minimum size of the meta-module was ≥ 3. Furthermore, meta-modules were summarized by their “eigenmodules”, defined as the first principal component of the module REI projections corresponding to each meta-module (“seed modules”). This is conceptually identical to summarizing gene co-expression modules by their eigengenes. Highly similar meta-modules were merged if their correlations exceeded 0.95 by combining the seed modules of each meta-module into a single meta-module. This pipeline was repeated for two regions: DFC (**Fig. 5b-d**) and MTG (**Fig. S8**).

In parallel, subclasses were clustered by applying a similar clustering strategy to the transpose of the previous analysis: subclass-by-subclass similarity matrices were produced by calculating pairwise Pearson correlations of all subclass REI projection indices over all modules. Like before, we used `pmin` to produce a consensus minimum subclass-by-subclass similarity matrix. This matrix was used as input into complete linkage hierarchical clustering via the `flashclust` package.

To visualize meta-modules, heatmaps of eigenmodules calculated from projection data from either Jorstad et al.^30^ or Gabitto et al.^29^ were displayed with the consensus meta-module dendrogram overlaid on top of each heatmap (**Fig. 5b-d, Fig. S8**). As eigenmodules are calculated using principal component analysis (PCA), the values of eigenmodules are arbitrary units that range from -1 to 1. Subclasses (y-axis of the heatmaps) were clustered according to the consensus subclass dendrogram. For each node of the meta-module dendrogram above a certain arbitrary threshold (distance = 0.3), nodes were labeled with two summary statistics: the percentage of all modules represented by all leaves under a given node (“m”), and the percentage of all module genes represented by those modules (“g”). The same statistics for each individual leaf are displayed as barplots below the dendrograms.

### 4. dCoPA

In addition to the neurotypical snRNA-seq samples from Gabitto et al.^29^ and Liu et al.^49^, we ran CoPA on disease samples from the same datasets. Gabitto et al. consists of 75 donors with Alzheimer’s disease representing 2,224,098 nuclei, while Liu et al. consists of 51 donors with Alzheimer’s disease representing 527,907 nuclei. We also downloaded data from the brainSCOPE resource^55^ (https://brainscope.gersteinlab.org/), which profiles adult human DFC samples from the PsychENCODE Consortium^64^ (https://www.psychencode.org/). Of the studies aggregated by the brainSCOPE resource, we analyzed studies from the CommonMind Consortium^42^ (53 control donors representing 287,783 nuclei and 47 schizophrenia donors representing 214,234 nuclei) and SZBDMulti-Seq^65^ (24 control donors representing 173,570 nuclei and 24 schizophrenia donors representing 171,313 nuclei).

#### 4.1 dCoPA overview

We leveraged CoPA projections to find disease-specific gene expression dysregulation at module-level resolution. To achieve this, we designed an accompanying pipeline (called “dCoPA” or differential CoPA). dCoPA selects a group of candidate disease-related modules based on the application of a series of filtering criteria to CoPA projections that ensure that any discovered disease associations are significant, interpretable, and reproducible (**Fig. 6a**).

The first filtering step was to determine whether module projections were significantly different between case and control samples for a given subclass. To do this, we projected modules from the integrated bulk control dataset onto case and control samples in mean log space. For each module (n = 1,016) and subclass (n = 24), we calculated the Euclidean distance between the mean expression vectors of all module genes in cases and controls. We compared this distance to a distribution of randomly generated distances, created by randomly generating and projecting 10,000 modules for each real module (for a total of 10,000 times 1,016 modules) that were matched in size to the corresponding real module. Each distance was assigned an empirical p-value representing the proportion of random distances smaller than the real distance. P-values were then adjusted according to the Benjamini-Hochberg procedure.

In addition to significance, we also filtered modules according to the directionality of module genes. We selected significant modules where module topmodposbc genes were either uniformly up- or down-regulated in disease samples relative to control samples. Any module where directionality was not consistent (e.g., some module genes showing higher expression in disease samples with others showing lower expression) was filtered out. Our rationale for applying this additional filtering step was to maximize the interpretability of the final list of dCoPA-selected modules.

#### 4.2 Analysis of the rate of discovery of significant projections from random modules

To ensure that the significance of projections could not be spuriously found in the data, we generated 1,000 random module networks, each consisting of the same number of modules (n = 1,016) as the real module network generated from the integrated bulk control dataset. Random modules were generated by randomly sampling all genes (n = 18,913), where each module of each random network was matched in size to the corresponding module in the real network. For each random network, we calculated the percentage of modules marked significant by dCoPA (boxplots, **Fig. S9a-d**), and we calculated the average percentage of subclasses per module marked significant by dCoPA (boxplots, **Fig. S9e-h**). We calculated the same percentages for the real network (red dots, **Fig. S9**).

#### 4.3 Comparison of control-versus disease-derived dCoPA modules

To assess the reproducibility of dCoPA modules, we applied dCoPA to projections produced from two sets of modules (derived from either the integrated bulk control dataset or the AD ROSMAP dataset^53^) and two snRNA-seq datasets (Gabitto et al.^29^ and Liu et al.^49^) split by region (DFC and MTG), for a total of eight sets of candidate dCoPA modules (**Fig. 7a-b**).

To find which dCoPA modules were common to both datasets, we identified the intersection per subclass between dCoPA modules found from Gabitto et al. versus dCoPA modules found from Liu et al. (bottom two rows of **Fig. 7a-b**). Subclasses were matched between the two datasets based on the mapping strategy described previously. We did this separately for each set of modules and each region.

To compare dCoPA modules derived from the integrated bulk control dataset (“control-derived”) to those derived from the AD ROSMAP dataset (“AD-derived”), we first found common dCoPA modules between datasets as described above. We then summarized common dCoPA modules for each subclass by the union of their topmodposbc genes. Finally, we performed one-sided Fisher’s exact tests comparing control-derived genes to AD-derived genes for each subclass (**Fig. S11**).

To assess the overlap of dCoPA genes across subclasses, we first found common genes that overlapped between control-derived and AD-derived genes per subclass. To visualize the overlap of these genes by subclass, we used these genes as input for the `UpSet` function from the `ComplexHeatmap`^66^ package (v2.22.0). Combinations where one or fewer genes overlapped between subclasses were not plotted for readability.

We repeated all of the above analyses for SCZ-derived modules and projections. We applied dCoPA to projections produced from two sets of modules (derived from either the integrated bulk control dataset or the SCZ Brainseq Phase 1 dataset^43^) and two snRNA-seq datasets (CMC^42^ and SZBDMulti-Seq^65^ from the brainSCOPE resource^55^).

#### 4.4 Summary of dCoPA genes

We created summary tables for dCoPA genes by subclass. For each subclass, we performed enrichment analysis for dCoPA genes associated with that subclass using Gene Ontology gene sets obtained from the Molecular Signatures Database (MSigDB v7.4). Enrichment tests were performed by using a one-sided Fisher’s exact test using the base R `phyper` function. The top three gene sets by Benjamini-Hochberg adjusted p-values per subclass were shown.

dCoPA genes were annotated for association with Alzheimer’s disease and schizophrenia with a single pipeline (Claude Sonnet 5, run via Claude Code [https://www.anthropic.com/claude-code]), applied independently to each disease. The reproducible cross-dataset dCoPA gene lists (i.e., genes belonging to modules that were significantly and uniformly dysregulated in the same cell type[s]) were associated with disease via one of three tracks. First, genes present in a curated disease database (OMIM^67^, Open Targets Platform^68^, ClinVar^69^, and, for AD, AlzForum [https://www.alzforum.org]) were admitted directly. Second, for all remaining genes, NCBI E-utilities (https://www.ncbi.nlm.nih.gov/books/NBK25497/) deterministically retrieved the total disease hit count and the top five most relevant PubMed abstracts for “GENE“[TIAB] AND “<disease>“[TIAB] (2000–present); because abstracts and PMIDs were fetched directly from NCBI, citation hallucination was impossible by construction. Claude Sonnet 5 then classified a gene as disease-linked only if a retrieved abstract established a direct mechanistic link to one of the disease’s mechanism categories (13 for AD, 14 for SCZ), rejecting co-mentions, differential-expression-only findings, and gene-symbol homonyms, and citing only fetched PMIDs. Genes missed by the exact symbol were re-queried using NCBI gene aliases under a homonym-specificity guard. Third, three AD genes whose literature refers to a shared protein-family name (PPP3CB/calcineurin, SPTAN1/spectrin, and EDEM3) were preserved from prior manual curation; schizophrenia required none. Each gene was annotated with its total PubMed hit count and, for AD, its Open Targets association score (disease MONDO_0004975), then intersected with the significant and reproducible dCoPA gene set and annotated with its normal and disease module membership, associated neuronal subclass(es), and direction of module change (**Table S15**). The final results comprised 74 (DFC) and 215 (MTG) genes for AD and 51 (DFC) genes for schizophrenia.

#### 4.5 CoPA Cabana (R Shiny app)

We made CoPA and dCoPA results explorable by developing an interactive web application with Claude using the R Shiny framework (bslib UI, plotly interactive graphics), which was deployed on https://oldhamlab.shinyapps.io/copacabana. The application comprises four tabs. The CoPA tab enables browsing and gene-based searching for gene coexpression modules identified in our integrated control cohort (**Fig. 3**), displaying the same information portrayed in **Fig. 3b-f** for all modules. The dCoPA tab visualizes all dCoPA-implicated modules for all disease comparisons explored in **Figs. 7**-**8**. The OMICON tab allows users to project coexpression modules downloaded from OMICON (https://theomicon.ucsf.edu) onto a number of select snRNA-seq reference atlases. The Gene Projection tab reports per-gene cell-type expression across reference datasets.

## ACKNOWLEDGMENTS

This work was supported by NIH/NIMH R01MH113896, NIH/NCI R01CA244621, NIH/NIMH R01MH123156, and NIH/NCI R01CA292649 (M.C.O.). We thank Aditya Kshirsagar for coining the CoPA acronym and all Oldham lab members for helpful discussions. We acknowledge all tissue donors and their families, the participating brain banks, and the investigators who generated and made these data available for reuse. The results published here are based on data from the sources listed below.

## AD Knowledge Portal

### ROSMAP

Study data were provided by the Rush Alzheimer’s Disease Center, Rush University Medical Center, Chicago. Data collection was supported through funding by NIA grants P30AG10161 (ROS), R01AG15819 (ROSMAP; genomics and RNAseq), R01AG17917 (MAP), R01AG30146, R01AG36836 (RNAseq), U01AG46152 (ROSMAP AMP-AD), U01AG61356 (ROSMAP AMP-AD), P30AG072975, the Illinois Department of Public Health (ROSMAP), and the Translational Genomics Research Institute (genomic). Additional phenotypic data can be requested at https://www.radc.rush.edu. All ROSMAP participants enrolled without known dementia and agreed to detailed clinical evaluation and brain donation at death. Both studies were approved by the Institutional Review Board of Rush University Medical Center; each participant signed an informed consent, Anatomic Gift Act, and repository consent permitting their data to be repurposed.

### SEA-AD

Study data were generated from postmortem brain tissue obtained from the University of Washington BioRepository and Integrated Neuropathology (BRaIN) laboratory and Precision Neuropathology Core, which is supported by the NIH grants for the UW Alzheimer’s Disease Research Center (P50AG005136 and P30AG066509) and the Adult Changes in Thought Study (U01AG006781 and U19AG066567). This study was supported by NIA grant U19AG060909.

### Liu et al

Study data were generated from postmortem brain tissue provided by the Religious Orders Study and Rush Memory and Aging Project (ROSMAP) cohort at Rush Alzheimer’s Disease Center, Rush University Medical Center, Chicago. This work was supported in part by the Cure Alzheimer’s Fund, NIH grants AG058002, AG062377, NS110453, NS115064, AG062335, AG074003, NS127187, MH119509, HG008155 (M.K.), RF1AG062377, RF1 AG054321, RO1 AG054012 (L.-H.T.) and the NIH training grant GM087237 (to C.A.B.). All donors contributing to this resource are ROS or MAP participants; the ROSMAP consent and IRB statement above therefore applies.

### CommonMind Consortium

#### CMC and CMC_HBCC

Data were generated as part of the CommonMind Consortium supported by funding from Takeda Pharmaceuticals Company Limited, F. Hoffmann-La Roche Ltd and NIH grants R01MH085542, R01MH093725, P50MH066392, P50MH080405, R01MH097276, R01MH075916, P50MH096891, P50MH084053S1, R37MH057881, AG02219, AG05138, MH06692, R01MH110921, R01MH109677, R01MH109897, U01MH103392, and contract HHSN271201300031C through IRP NIMH. Brain tissue for the study was obtained from the following brain bank collections: the Mount Sinai NIH Brain and Tissue Repository, the University of Pennsylvania Alzheimer’s Disease Core Center, the University of Pittsburgh NeuroBioBank and Brain and Tissue Repositories, and the NIMH Human Brain Collection Core. CMC data have since migrated to the NIMH Data Archive (https://nda.nih.gov/).

### PsychENCODE Consortium

#### BrainGVEX, BrainSeq Phase 1, CMC_HBCC, and the Capstone II harmonized data / brainSCOPE

Data were generated as part of the PsychENCODE Consortium, supported by:

U01DA048279, U01MH103339, U01MH103340, U01MH103346, U01MH103365, U01MH103392, U01MH116438, U01MH116441, U01MH116442, U01MH116488, U01MH116489, U01MH116492, U01MH122590, U01MH122591, U01MH122592, U01MH122849, R01MH094714, R01MH105472, R01MH105898, R01MH109677, R01MH109715, R01MH110905, R01MH110920, R01MH110921, R01MH110926, R01MH110927, R01MH110928, R01MH111721, R01MH117291, R01MH117292, R01MH117293, R21MH102791, R21MH103877, R21MH105853, R21MH105881, R21MH109956, R56MH114899, R56MH114901, R56MH114911, and P50MH106934.

PsychENCODE data have since migrated to the NIMH Data Archive (https://nda.nih.gov/pec; NDA collection 5032).

### BrainSeq Consortium

Data were generated by the BrainSeq Consortium at the Lieber Institute for Brain Development.

### Genotype-Tissue Expression (GTEx) Project

The Genotype-Tissue Expression (GTEx) Project was supported by the Common Fund of the Office of the Director of the National Institutes of Health, and by NCI, NHGRI, NHLBI, NIDA, NIMH, and NINDS. The data used for the analyses described in this manuscript were obtained from dbGaP accession number phs000424.v8.p2 on 20 April 2023. These data were accessed through the NHGRI AnVIL Project; the authors wish to acknowledge the NHGRI AnVIL Project (Schatz, M.C. et al. Cell Genomics 2, 100085 (2022); https://anvilproject.org).

### NABEC

NABEC data were accessed from dbGaP (https://dbgap.ncbi.nlm.nih.gov/home/); the study was supported in part by the Intramural Research Program of the National Institute on Aging, NIH (project Z01 AG000949).

### BICCN (Jorstad et al.)

The data analyzed in this study were produced through the Brain Initiative Cell Census Network (BICCN:RRID:SCR_015820) and deposited in the NeMO Archive (RRID:SCR_002001) under identifiers nemo:dat-rg2rc5m (https://assets.nemoarchive.org/dat-rg2rc5m) and nemo:dat-swzf4kc (https://assets.nemoarchive.org/dat-swzf4kc).

### Morabito et al

These data are publicly available through the Gene Expression Omnibus (GSE174367) and are acknowledged by citation.

## AUTHOR CONTRIBUTIONS

G.K. and M.C.O. conceptualized analyses, planned analyses, interpreted results, and wrote the manuscript. G.K. performed all analyses, produced all figures and tables, and created the CoPA Cabana web site.

## DATA AND CODE AVAILABILITY

All analyses are based on published datasets accessed as described in Methods. Results of our analyses are available in supplementary tables and on the CoPA Cabana web site that accompanies this study (https://oldhamlab.shinyapps.io/copacabana/). Accession identifiers and repositories for all datasets are provided in Table S1 (snRNA-seq) and Table S8 (bulk RNA-seq). Code to reproduce our analyses is available in the Oldham Lab GitHub repository (https://github.com/oldham-lab/kang-oldham-2026).

## EXTENDED DATA FIGURE LEGENDS

**Fig. S1.**
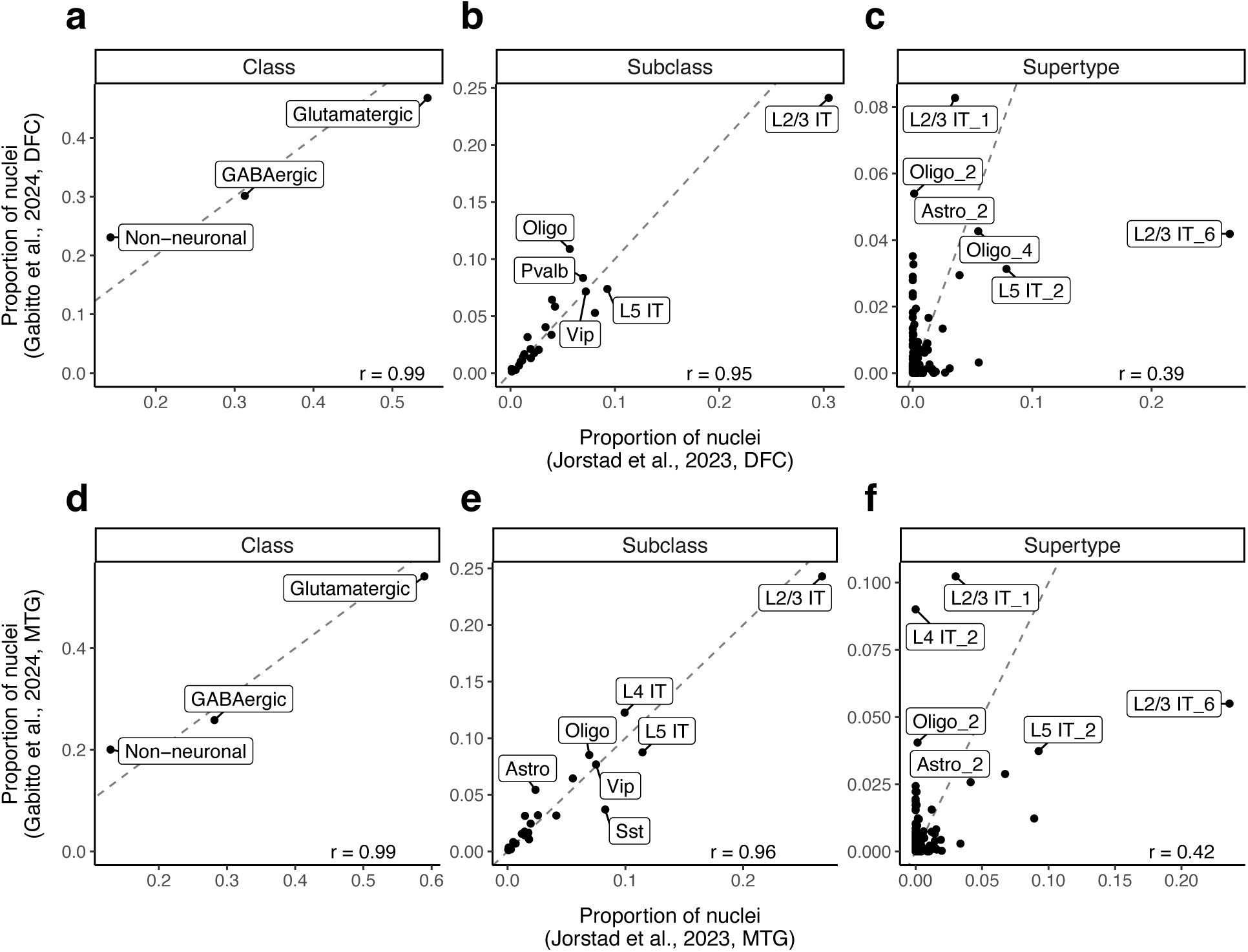
Cell class and subclass proportions are more highly conserved than supertype proportions in human neocortex. **a-f)** Proportion of nuclei in adult human dorsal prefrontal cortex (DFC: **a-c**) or middle temporal gyrus (MTG: **d-f**) assigned to each cell class (**a, d**), subclass (**b, e**), or supertype (**c, f**) in two snRNA-seq datasets^29,30^.

**Fig. S2.**
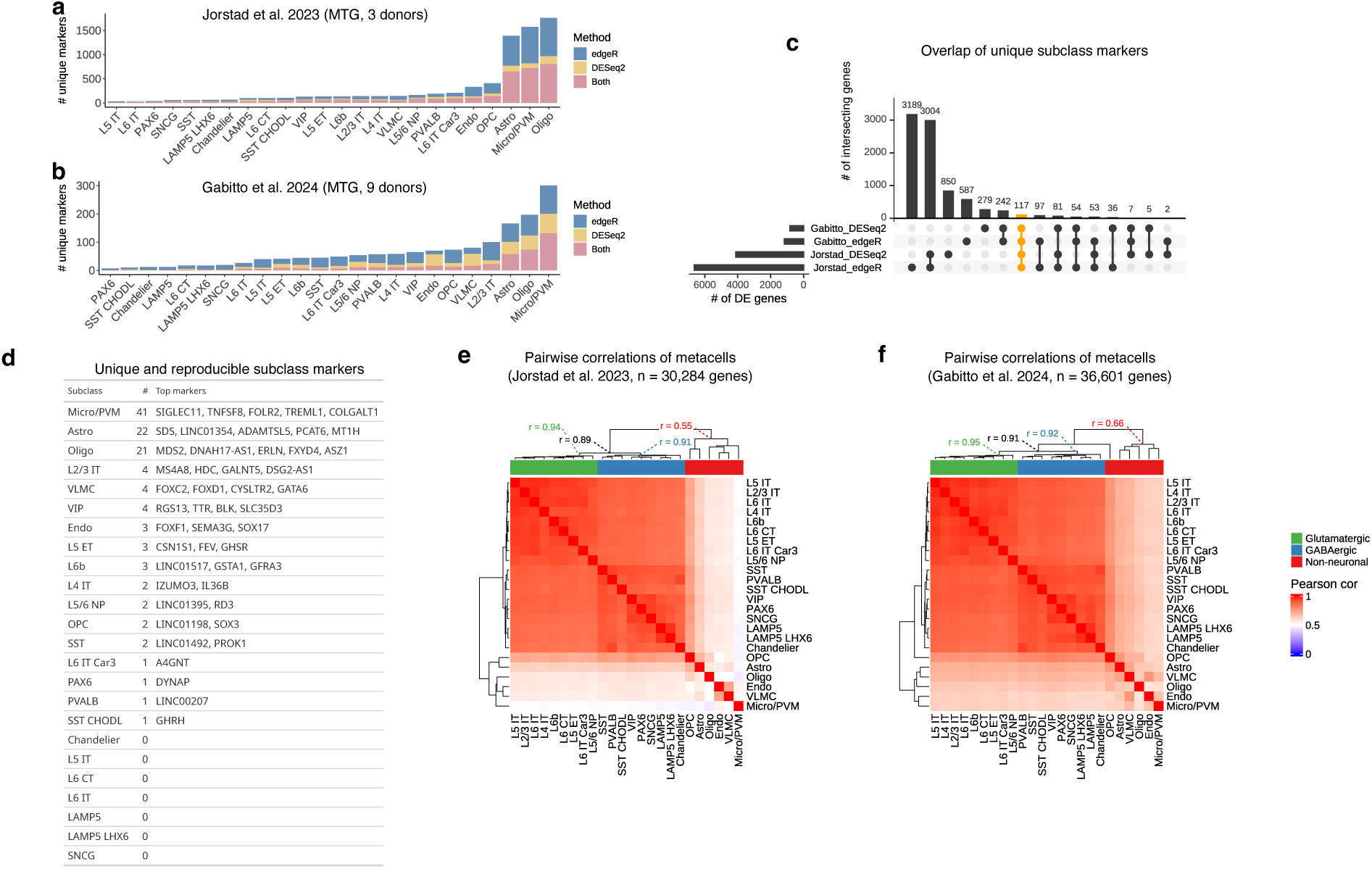
Cell type marker genes in human temporal cortex vary by choice of study and algorithm. **a-b)** Number of unique marker genes (FDR < .05) identified by differential expression (DE) analysis of snRNA-seq data from two studies^29,30^ of normal adult human middle temporal gyrus (MTG) using `edgeR`^33^, `DESeq2`^34^, or both. **c)** Overlap of unique marker genes from **a** & **b** for all cell types. **d**) Unique and reproducible marker genes for each cell type ranked by significance. **e-f**) Heatmaps of genome-wide correlations among pseudobulked cell types from Jorstad et al.^30^ (**e**) and Gabitto et al^29^. (**f**). Mean correlations among metacells of a given class (glutamatergic, GABAergic, neuronal, non-neuronal) are reported at dendrogram branch points.

**Fig. S3.**
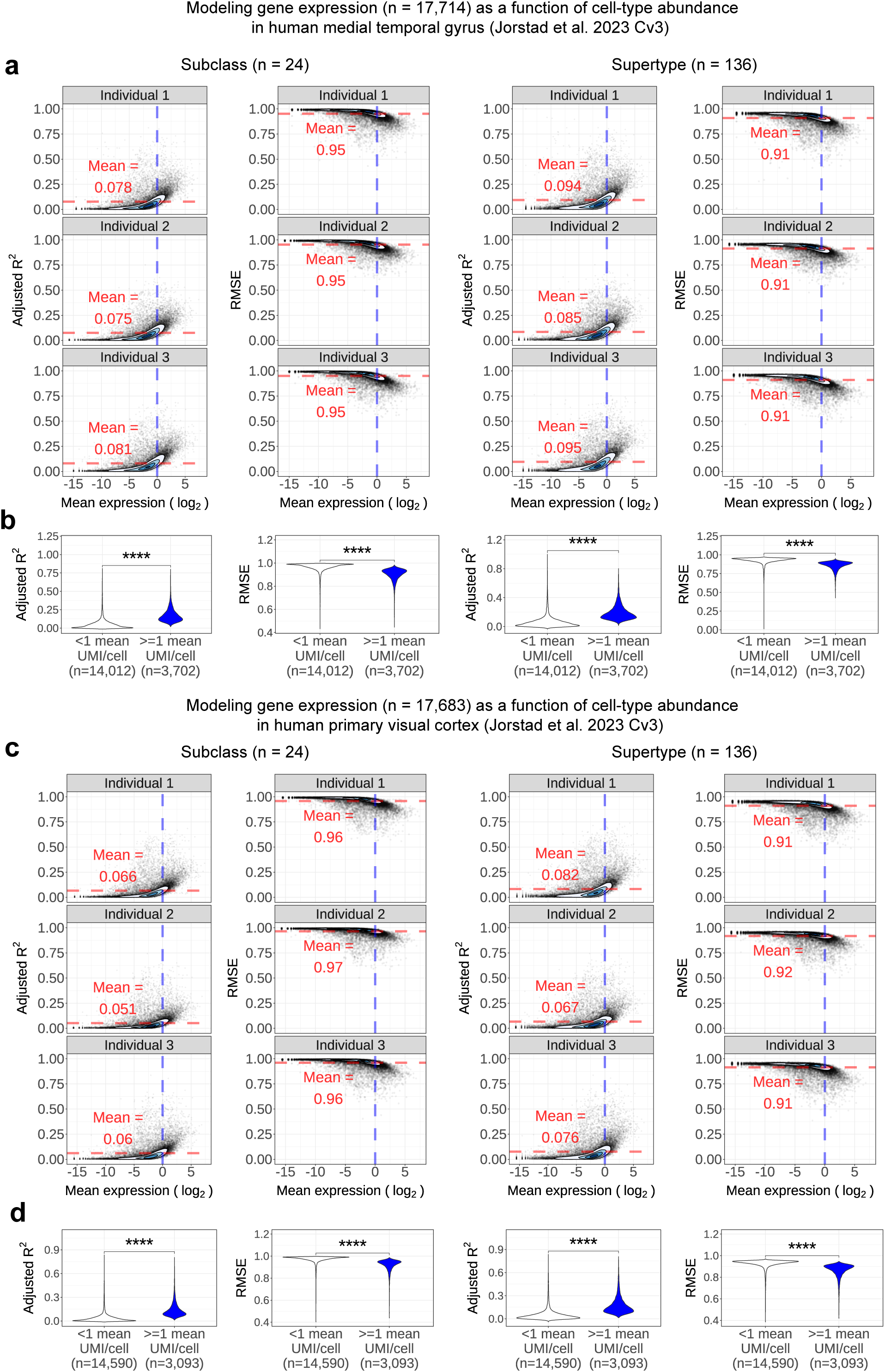
Genome-wide expression variation in pseudobulked snRNA-seq data from human temporal and visual cortex is poorly explained by cell-type abundance, particularly for low-expressed genes. **a-d**) Variation in cell type abundance was used to predict genome-wide expression levels in pseudobulked snRNA-seq data^30^ from middle temporal gyrus (**a**, **b**) or primary visual cortex (**c**, **d**) via multiple linear regression as described in Fig. 2. (**a**, **c**) Adjusted R^2^ and root mean square error (RMSE) modeling results for all genes vs. mean expression level (log_2_). Red dotted lines denote mean modeling performance, while blue dotted lines denote mean expression of 1 UMI/nucleus^19,20^. (**b**, **d**) Violin plots of modeling performance from (**a**, **c**), stratified by genes below and above this critical threshold. ****P < 0.0001

**Fig. S4.**
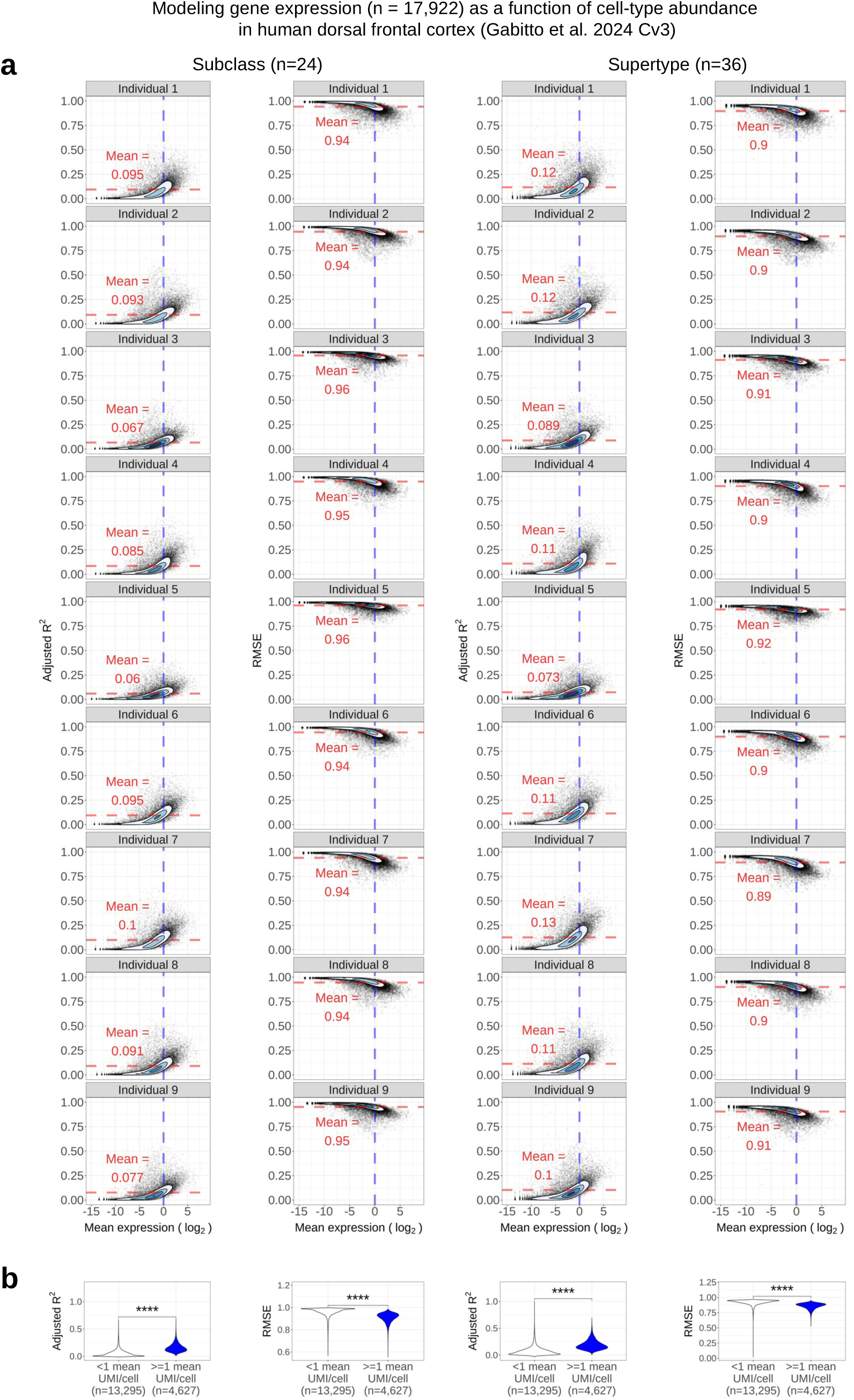
Genome-wide expression variation in pseudobulked snRNA-seq data from human frontal cortex is poorly explained by cell-type abundance, particularly for low-expressed genes. **a-b)** Variation in cell type abundance was used to predict genome-wide expression levels in pseudobulked snRNA-seq data^29^ from dorsal frontal cortex via multiple linear regression as described in Fig. 2**.a**) Adjusted R^2^ and root mean square error (RMSE) modeling results for all genes vs. mean expression level (log_2_). Red dotted lines denote mean modeling performance, while blue dotted lines denote mean expression of 1 UMI/nucleus^19,20^. (**b**) Violin plots of modeling performance from (**a**), stratified by genes below and above this critical threshold. ****P < 0.0001

**Fig. S5.**
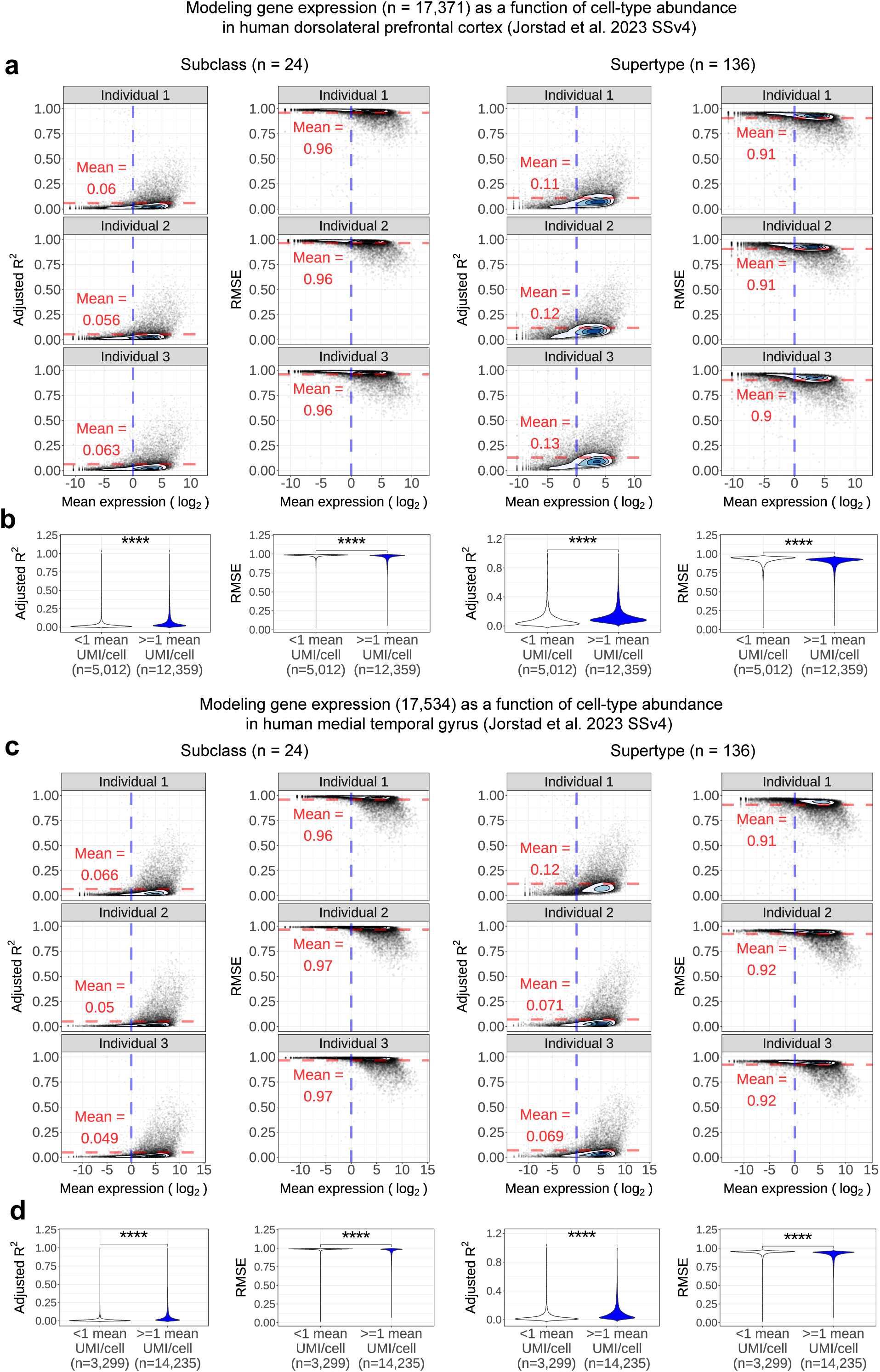
Genome-wide expression variation in pseudobulked snRNA-seq data from human neocortex is poorly explained by cell-type abundance, even with deeper sequencing. **a-d**) Variation in cell type abundance was used to predict genome-wide expression levels in pseudobulked snRNA-seq data^30^ from frontal cortex (**a**, **b**) or temporal cortex (**c**, **d**) via multiple linear regression as described in Fig. 2. (**a**, **c**) Adjusted R^2^ and root mean square error (RMSE) modeling results for all genes vs. mean expression level (log_2_). Red dotted lines denote mean modeling performance, while blue dotted lines denote mean expression of 1 UMI/nucleus^19,20^. (**b**, **d**) Violin plots of modeling performance from (**a**, **c**), stratified by genes below and above this critical threshold. ****P < 0.0001

**Fig. S6.**
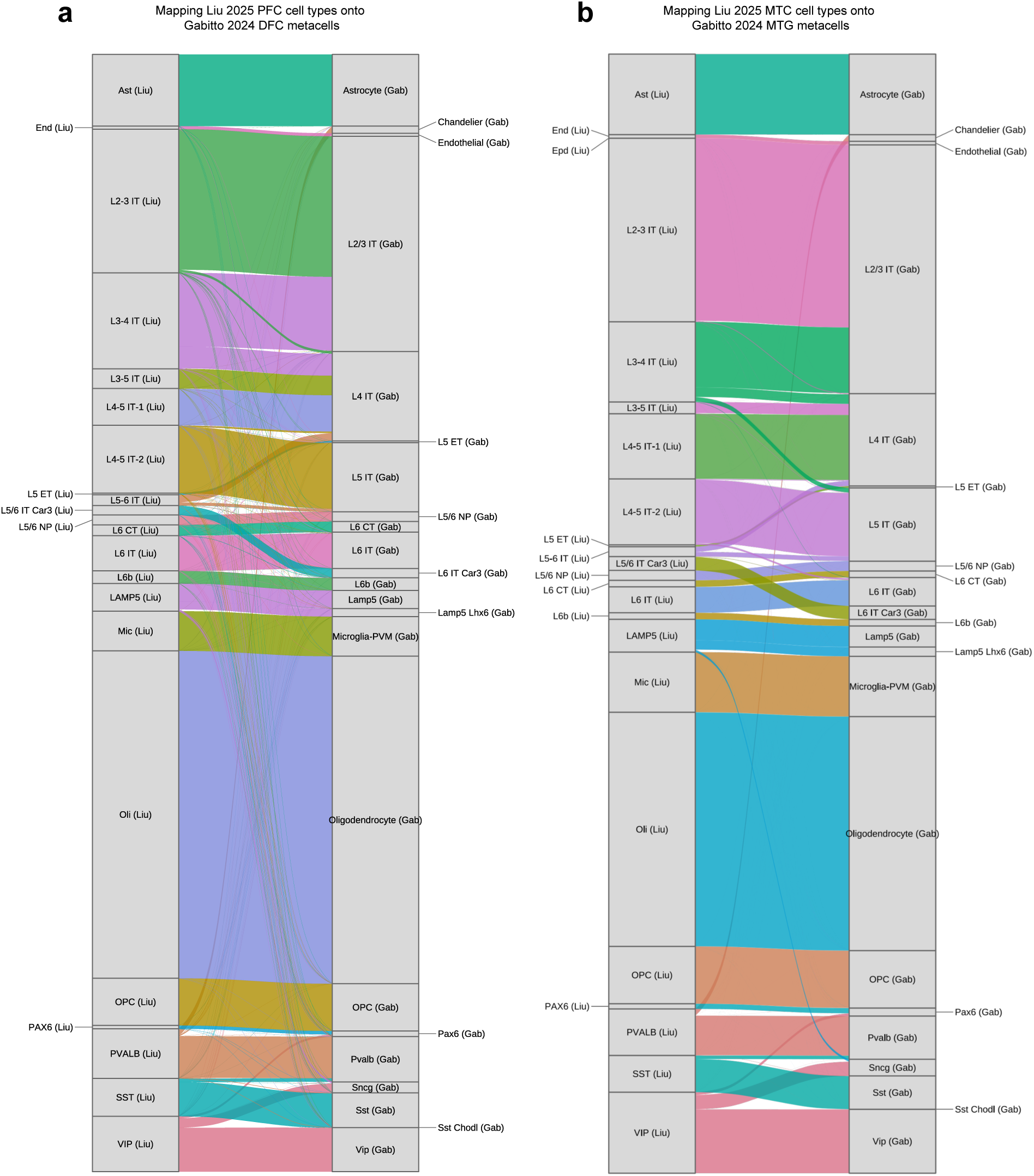
Liu et al. nuclei map cleanly to subclass identities from Gabitto et al. **a-b)** River plots display mapping of human frontal cortex (**a**) and temporal cortex (**b**) nuclei from Liu et al.^49^ to subclass identities from Gabitto et al^29^.

**Fig. S7.**
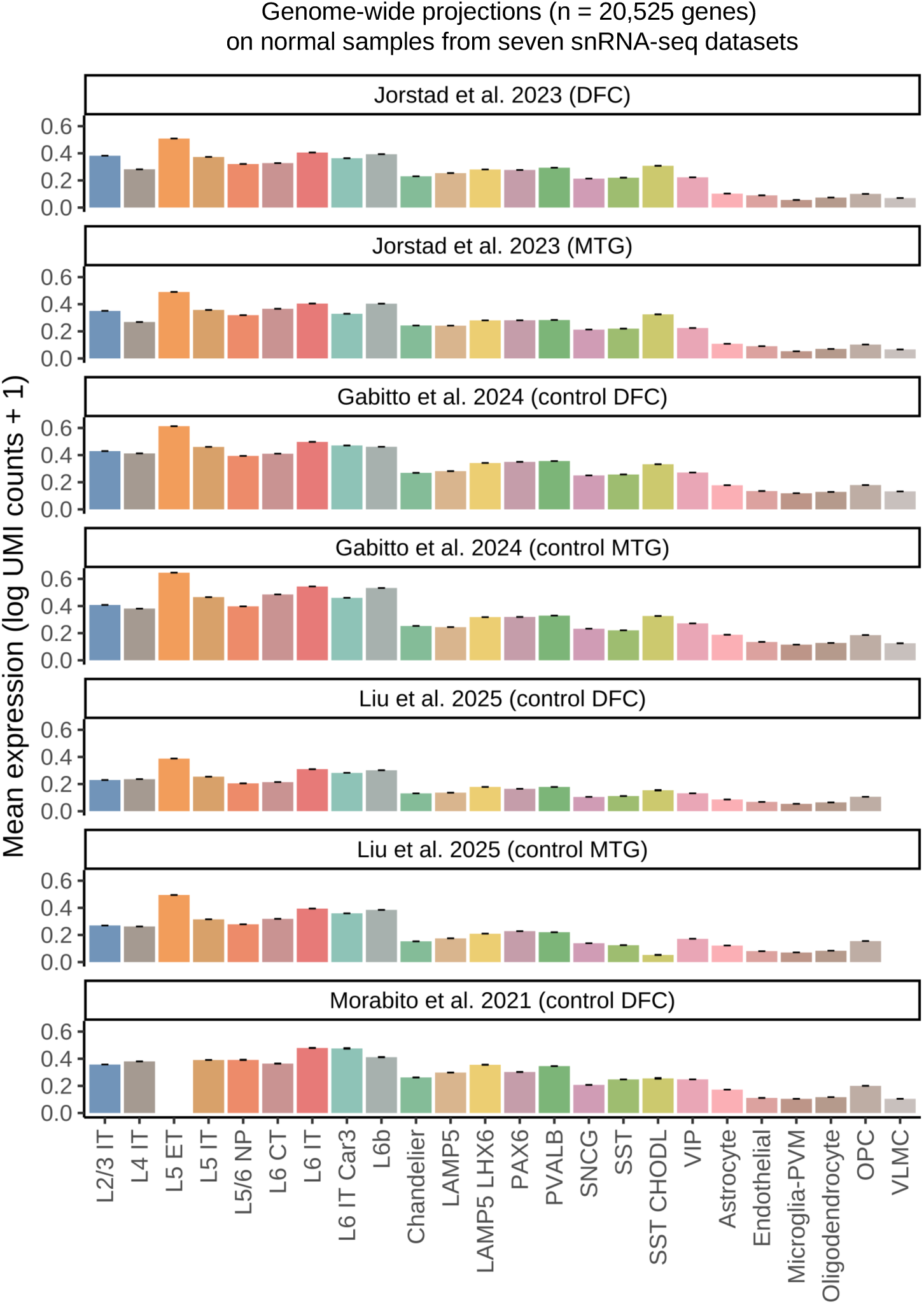
Genome-wide expression levels vary significantly among human neocortical subclasses. Expression levels for all genes (n = 18,913) were averaged for each cell type using snRNA-seq data from four studies^29–31,49^ and two brain regions. DFC = dorsal frontal cortex; MTG = middle temporal gyrus.

**Fig. S8.**
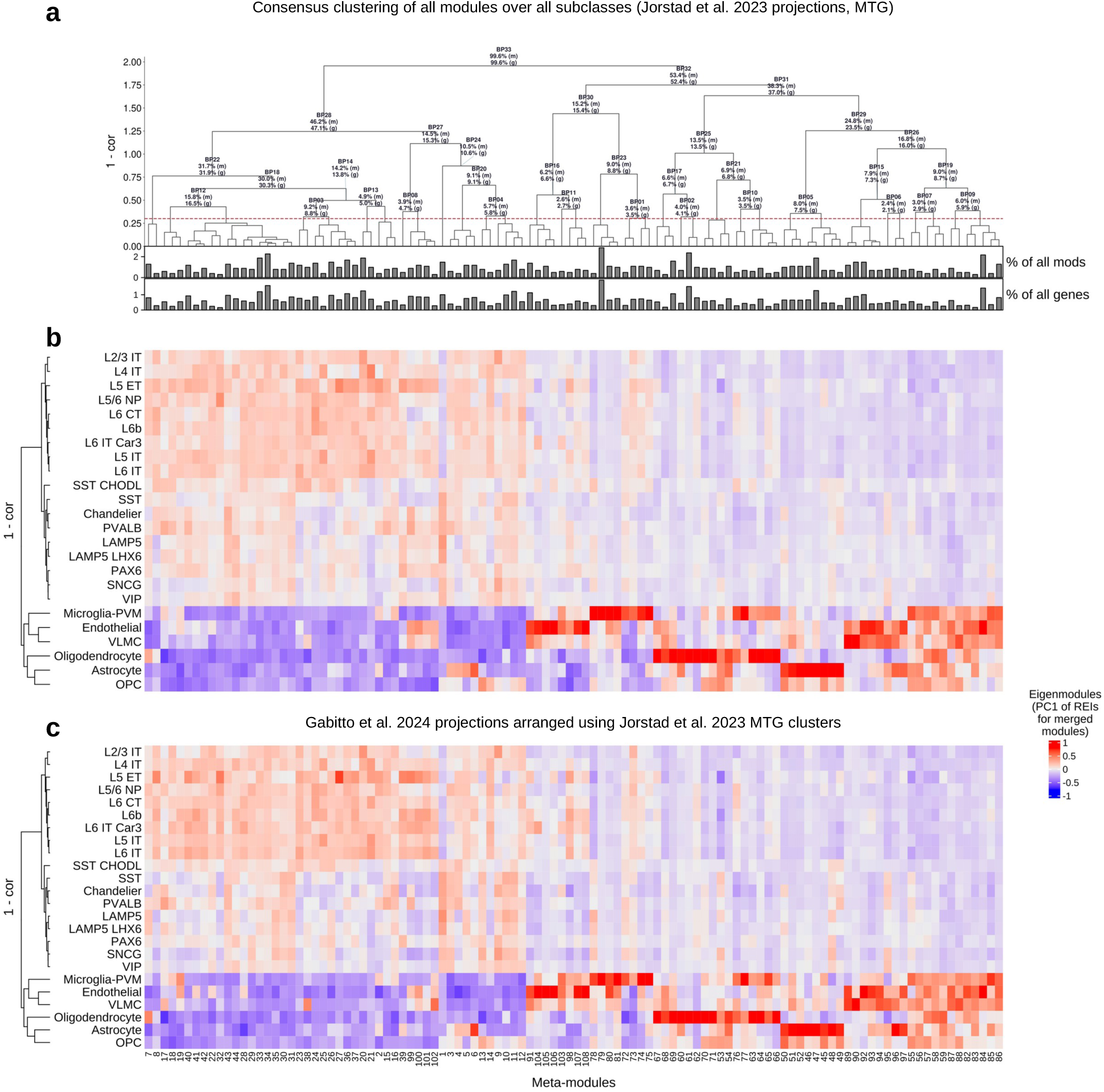
Clustering CoPA projection patterns reveals the major motifs of gene activity in human middle temporal gyrus (MTG) cell types. Clustering was performed as described in Fig. 5a. **a**) Hierarchical clustering of meta-modules (n=108) using MTG snRNA-seq data from Jorstad et al^30^. Barplots below the dendrogram report the percentages of all modules (top) and genes (bottom) comprising each meta-module. Branch points above the red dashed line report the percentages of all constituent modules (‘m’) and genes (‘g’). **b-c**) Heatmaps of eigenmodules (first principal components of REI vectors for merged modules) produced using MTG snRNA-seq data from Jorstad et al.^30^ (**b**) and Gabitto et al.^29^ (**c**). Cell types were clustered as described in Fig. 5a.

**Fig. S9.**
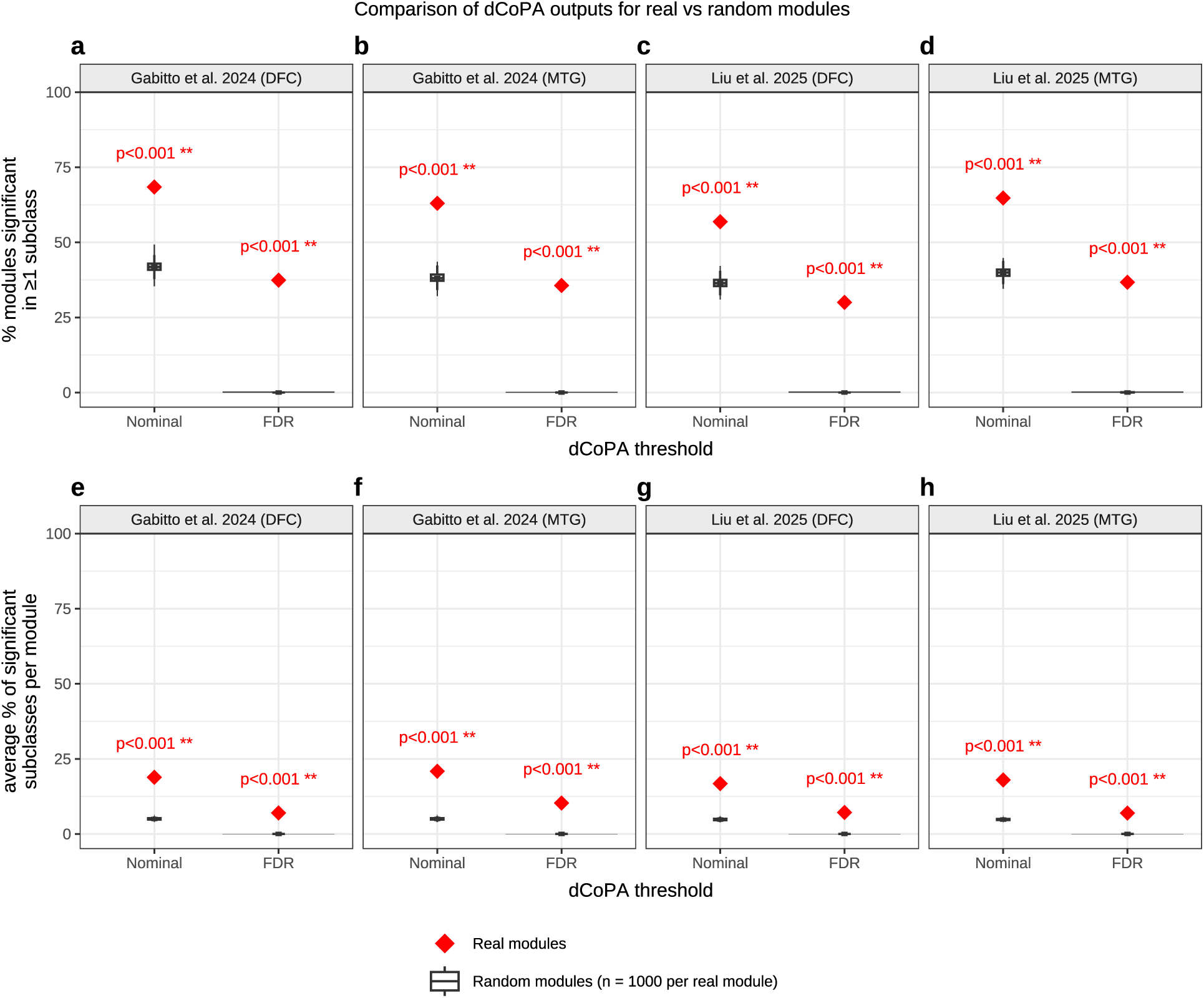
Application of dCoPA to random modules produces no significant results. Comparison of dCoPA results for real modules (n=1,016) and random modules (formed by randomly selecting genes to match real module sizes [n=1,000 random modules per real module]). Four snRNA-seq datasets were analyzed from two studies^29,49^ and two brain regions: dorsal frontal cortex (DFC) and middle temporal gyrus (MTG). **a-d)** Percentage of all real (red) or random (black) modules with at least one significant cell type difference. **e-h)** The average percentage of significant subclasses per module. After correcting for multiple comparisons (FDR), no significant cell type differences were identified in random modules.

**Fig. S10.**
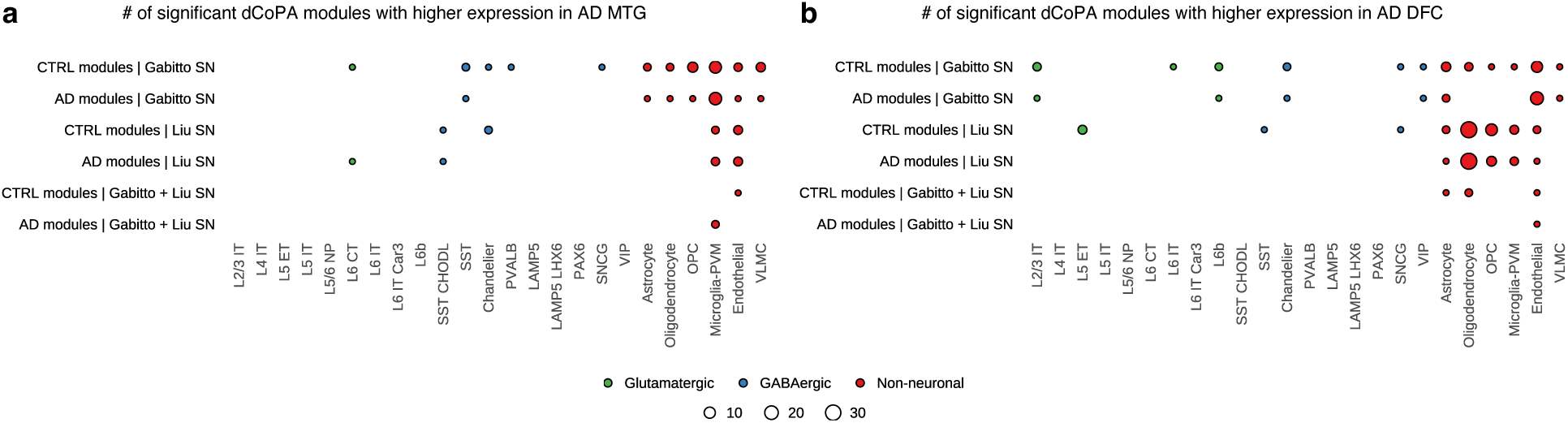
Up-regulation of modular gene activity in Alzheimer’s disease (AD) occurs mostly in non-neuronal cells. **a-b)** Dotplots depict the number of bulk coexpression modules identified by dCoPA that were significantly up-regulated in AD vs. CTRL cell types from MTG (**a**) or DFC (**b**) in snRNA-seq data from Gabitto et al.^29^, Liu et al^49^, or both (gold rows). CTRL / AD modules denote bulk coexpression modules derived from CTRL or AD samples, respectively. Interactive dot plots (**a-b**) are available on the CoPA Cabana web site (https://oldhamlab.shinyapps.io/copacabana/).

**Fig. S11.**
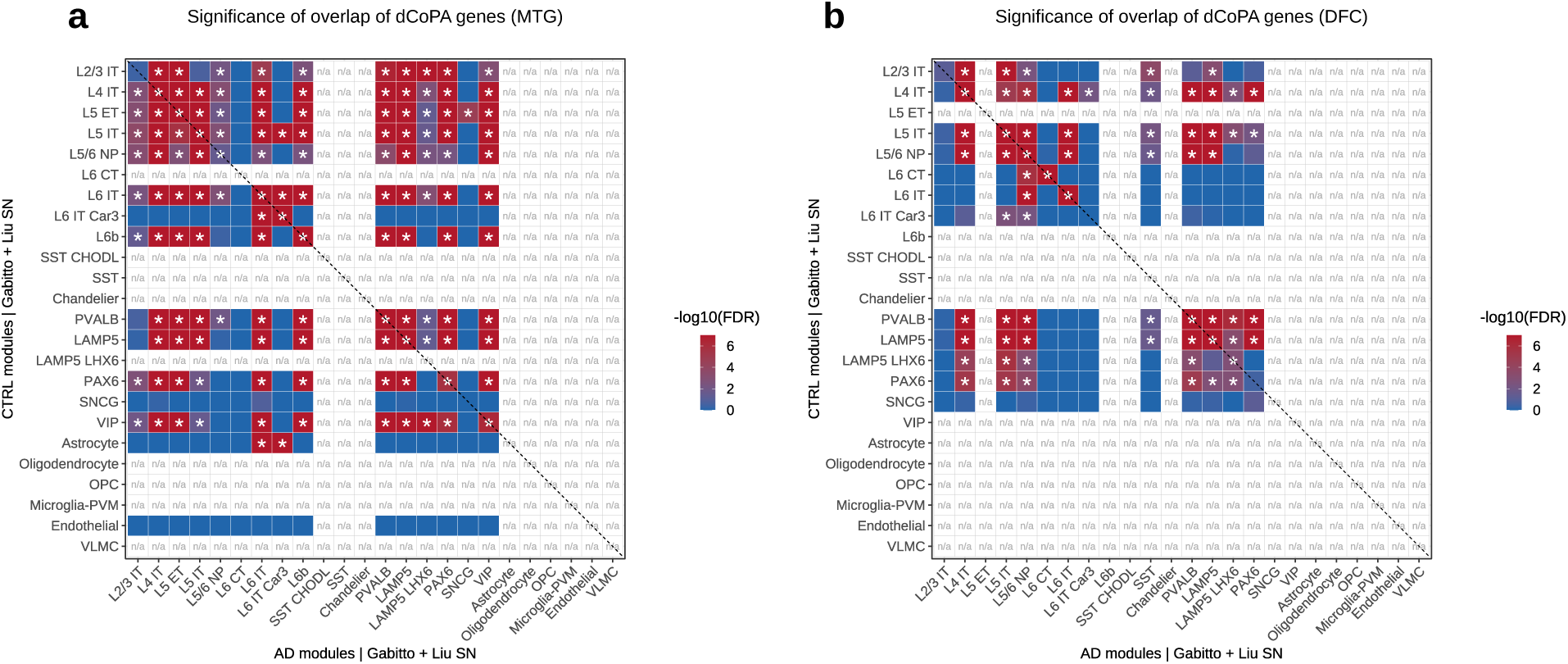
Modular gene activity is reproducibly and coordinately down-regulated in specific neuronal subclasses. **a-b)** Significance of overlap (one-sided Fisher’s exact test) for shared dCoPA genes with lower expression in AD MTG (a) and DFC (b) (i.e., the intersection of all genes for each subclass in the bottom two rows in Fig. 7a (**a**) and Fig. 7b (**b**).

## EXTENDED DATA TABLE LEGENDS

**Table S1 | snRNA-seq datasets analyzed in this study**

**Table S2 | Differential expression analysis of dorsal frontal cortex subclasses from Jorstad et al**^30^.

**Table S3 | Differential expression analysis of dorsal frontal cortex subclasses from Gabitto et al**^29^.

**Table S4 | Differential expression analysis of middle temporal gyrus subclasses from Jorstad et al**^30^.

**Table S5 | Differential expression analysis of middle temporal gyrus subclasses from Gabitto et al**^29^.

**Table S6 | Pseudobulk modeling of gene expression as a function of cell-type abundance (Jorstad et al**^30^)

**Table S7 | Pseudobulk modeling of gene expression as a function of cell-type abundance (Gabitto et al.**^29^)

**Table S8 | Bulk RNAseq datasets analyzed in this study**

**Table S9 | Assignments of all genes to all modules identified by CoPA in normal human frontal cortex**

**Table S10 | REI values for all modules and meta-modules in human neocortex.**

**Table S11 | Differential expression analysis of control vs. Alzheimer’s disease subclasses**

**Table S12 | Assignments of all genes to all modules identified by CoPA in frontal cortex from patients with Alzheimer’s disease**

**Table S13 | Significant dCoPA modules by cell type Table S14 | Significant dCoPA genes by cell type**

**Table S15 | AI-powered literature review for significant and reproducible dCoPA genes**

**Table S16 | Assignments of all genes to all modules identified by CoPA in frontal cortex from patients with schizophrenia**

